# Single-cell-scale spatial transcriptome reveals early regional priming of the developing mouse ovary

**DOI:** 10.1101/2025.09.16.676655

**Authors:** Anthony S. Martinez, Tyler J. Gibson, Courtney Diamond, Jennifer Jaime, Jennifer McKey

## Abstract

Mammalian ovary development is essential for female fertility, involving the complex spatial patterning of diverse cell types to establish the finite reserve of ovarian follicles. To uncover molecular mechanisms driving regionalization of the ovary while maintaining its native cellular architecture, we used Visium HD spatial transcriptomics. Our study captures all ovarian cell types at eight timepoints, generating a near single cell resolution library of spatial gene expression across development. Our analysis identified a previously uncharacterized "medullary core" domain that emerges by E16.5. Pre-granulosa cells (PGs) within this core express genes typically restricted to growing follicles in the postnatal ovary (*Nr5a2*, *Slc18a2*, *Ephx2)* suggesting core PGs are poised for activation while still in the fetal environment. This study provides a comprehensive architectural and molecular blueprint of the ovary, representing a fundamental resource for the investigation of regulatory mechanisms driving spatial patterning of the ovary and opens new avenues to explore the spatial determinants of female fertility and reproductive longevity.

**Teaser:** Visium HD provides a comprehensive blueprint of the mouse ovary, revealing a medullary core of activation-poised pregranulosa cells.

## Introduction

The ovarian reserve, the finite pool of follicles responsible for the entire reproductive lifespan of the adult mammalian female, is established during fetal and perinatal development (*1*, *2*). Thus, differences in ovary development have lifelong impacts on female health and fertility. A smaller reserve can lead to primary ovarian insufficiency (POI), characterized by subfertility and systemic health risks associated with early menopause (*3*). However, the mechanisms governing the highly dynamic and complex process of ovary development remain incompletely understood.

During fetal development, the gonad first arises as a bipotential primordium that is colonized by migrating primordial germ cells (*1*–*6*). At the stage of sex determination, around embryonic day (E)11.5 in mice, gonads in XX embryos commit to a female fate and develop into ovaries (*6*, *10*, *11*). This coincides with mitotic proliferation of germ cells that undergo incomplete cytokinesis to form germline cysts (*12*–*14*), and differentiation of bipotential supporting cells into pre-granulosa cells (PGs) (*6*, *15*–*18*). By E14.5 germline cysts are surrounded by PGs, forming transient structures termed ovigerous cords anchored in the center of the ovary (*19*, *20*). Around E17.5, ovigerous cords begin to break down to form individual ovarian follicles defined by the assembly of one oocyte and 2-3 pre-granulosa cells (*21*–*24*). These first follicles are assembled in the ovarian medulla with PGs arising from bipotential progenitors of the coelomic epithelium (CE) prior to sex determination (*24*, *25*). A second wave of PGs derive from the ovarian surface epithelium (OSE) after sex determination and assemble with oocytes closer to the surface to form cortical follicles (*24*, *26*–*29*). The regionalization of follicles into cortical and medullary pools holds functional significance, as from postnatal (P) day 2 to P7, follicles located in the medulla activate and begin to synchronously grow in a process termed first wave folliculogenesis (*18*, *30*–*32*). These perinatally-activated medullary follicles are largely depleted by the onset of puberty and only represent a small contribution to female fertility (*26*, *31*, *33*). The depletion of medullary follicles leaves a significantly larger pool of quiescent follicles in the cortex that constitute the ovarian reserve responsible for female fertility. It is from this pool, at the onset of puberty, that cyclical recruitment and ovulation of oocytes will occur (*24*, *31*, *32*). Because this reserve is finite, the mechanisms governing follicle regionalization and the rate of cyclical reduction are major determinants of female reproductive longevity.

The specification of somatic lineages from distinct progenitor sources is critical to the regional architecture of the ovary. While CE-derived PGs settle in the medulla and OSE-derived PGs populate the cortex, a third lineage of supporting-like cells (SLCs) emerges on the dorsal side of the gonad as early as E10.5 (*34*). These SLCs represent a progenitor population that gives rise to both the epithelial network of the rete ovarii and a subset of medullary PGs (*35*). The physical folding of the ovary between E14.5 and E16.5 internalizes these dorsal structures, creating a more heterogenous medullary microenvironment where the rete ovarii, SLCs and CE derived PGs are in close proximity (*29*). However, whether the local signaling milieu of the medulla or the intrinsic lineage of these cells dictates their eventual entry into the active follicle program during first wave folliculogenesis remains poorly defined. Recent studies have challenged the view that medullary follicles are a “sacrificed” population by identifying a correlative link between their activation and the expression of steroidogenic machinery in granulosa cells, which may contribute to the hormonal milieu of the juvenile female (*26*). While these first wave follicles were initially considered to only contribute to the first litter in mice (*31*), lineage-tracing indicates that a subset of follicles activated in perinatal life can stall their growth and contribute offspring throughout the fertile lifespan in mice (*33*). These data suggest that first-wave follicles follow divergent developmental trajectories. However, how these functional subsets emerge, and whether they represent a spatially segregated sub-compartment within the medulla or are stochastically distributed remains to be determined.

Recent single cell/nucleus transcriptomic (sc/snRNAseq) studies have allowed better appreciation of the temporal dynamics of gene expression networks during differentiation of ovarian cell types in mice (*2*, *15*, *25*, *34*, *36*–*41*). While these methodologies have revealed transcriptional signatures and gene regulatory networks underlying ovarian development, the dissociation of tissue required for these methods disrupts the spatial context necessary to understand the links between spatial patterning and the molecular specification of ovarian cell types. In addition, in traditional scRNA-seq datasets, the transcriptomic signatures of small, localized sub-compartments are often lost to the averaging effects of molecular clustering. In fact, recent 3D-imaging studies of ovary morphogenesis have highlighted the spatially dynamic nature of ovarian differentiation and establishment of the ovarian reserve (*10*, *29*, *32*, *42*–*45*). For example, we now know that meiotic entry of oocytes follows an anterior-posterior and radial pattern (*43*), that regionalization of the ovary into cortex and medulla coincides with physical folding of the ovarian domain (*29*), and that medullary follicles activate first in the dorsal region of the ovary during first wave folliculogenesis (*32*). However, we lack a high-resolution molecular map that correlates these morphogenic movements with transcriptional output.

To resolve how gene expression networks integrate spatial cues to establish regional patterns, we performed genome-wide spatial profiling of gene expression across the ovary throughout development. We used 10x Genomics Visium HD technology to generate a near single-cell resolution spatial transcriptomic time course of the mouse ovary across eight stages: four fetal (E12.5, E14.5, E16.5, and E18.5) and four postnatal (P0, P3, P21, and P90). These stages encompass the complete ontogeny of the ovary, from initial specification and physical folding to the establishment of the quiescent ovarian reserve and adult homeostasis. This approach identified a previously uncharacterized medullary core domain that emerges by E16.5. We demonstrate that PGs within this core possess an "activation-poised" molecular signature normally restricted to postnatal growing follicles, suggesting that the determinants of first-wave folliculogenesis are established well before birth. This study provides a comprehensive architectural and molecular blueprint of the ovary, offering new insights into the developmental origins of the ovarian reserve and the etiology of primary ovarian insufficiency.

## Results

### Multi-orientation embedding strategy captures a highly resolved time course across ovarian development and maturation

To generate a comprehensive dataset resolving ovarian development and maturation in the wild type CD-1 mouse, we selected four fetal stages (E12.4, E14.5, E16.5 and E18.5) and four postnatal stages (P0, P3, P21, and P90). We maximized use of the Visium HD slide (10X Genomics, USA) area by utilizing a high density, multi-orientation embedding strategy (**Fig. 1A**). For each timepoint, multiple ovaries from at least two litters were embedded on either their dorsal or ventral side in a single cryomold. Blocks were then sectioned, stained, and optimal sections were selected for sequencing (**Fig. S1**). This approach produced one section per stage containing several ovarian profiles, each of which were sectioned at different planes in the dorsal-ventral axis (**Fig. 1B**, **C**). The inclusion of peripheral tissues during embedding, namely the mesonephros (fetal only), the oviduct, the uterus, and the rete ovarii, further allowed for the preservation of key contextual structures.

**Figure 1.**
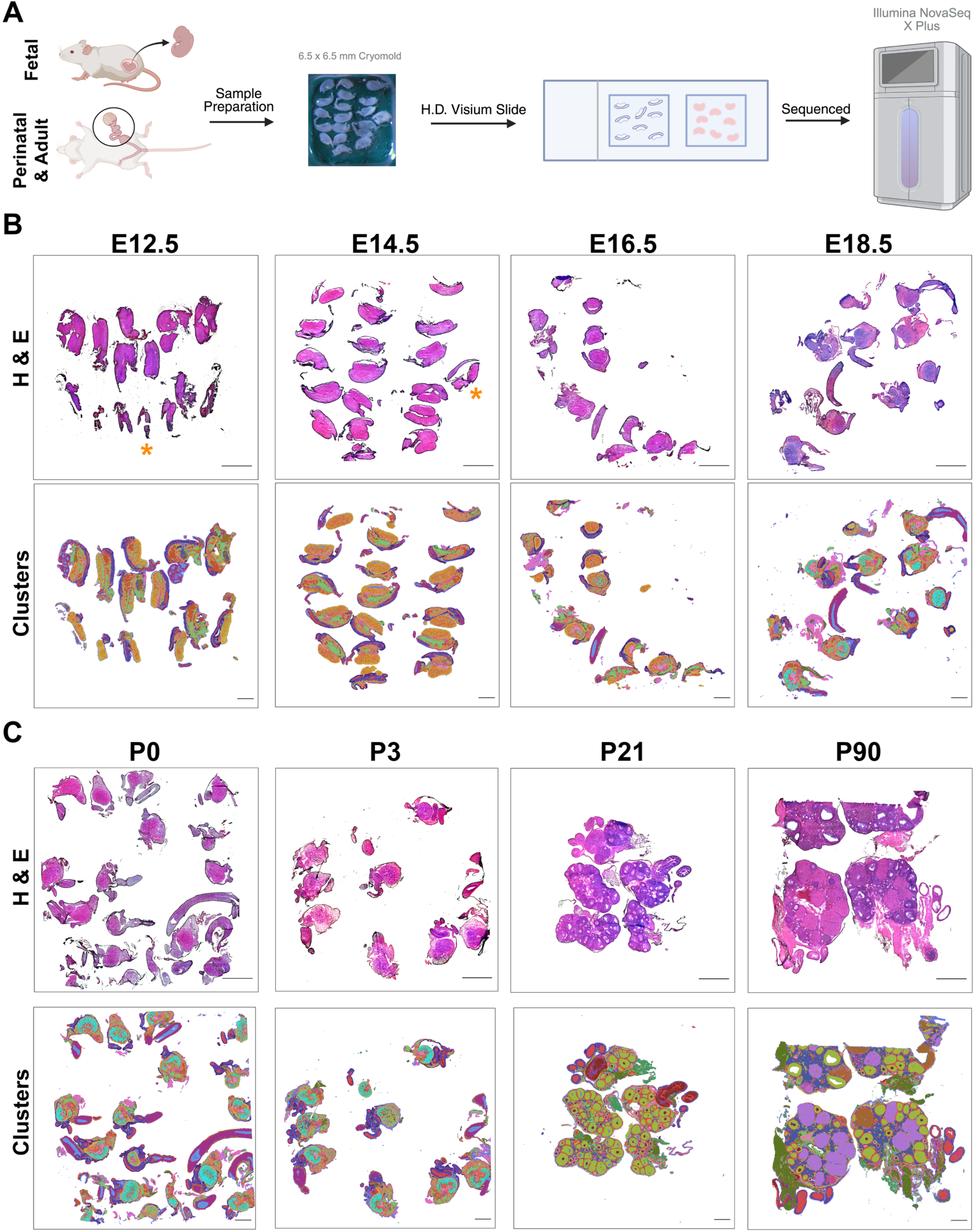
Multi-orientation embedding strategy allows robust capture of mouse ovarian development. (**A**) Overview of workflow: fetal and postnatal ovaries were dissected and embedded in a 7 x 7 cryomold. Optimal 10 µm sections were selected and sequenced. Created in BioRender. Martinez, A. (2025) https://BioRender.com/sytaju5 (**B-C**) *Top row*: 20x high resolution histology images of ovaries stained with H&E prior to sequencing. Scale bars, 1000 μm. *Bottom row*: Projection of 34 clusters identified in the integrated spatial dataset projected onto the tissue for each timepoint. Scale bars, 1000 μm. (**B**) Fetal stages: E12.5, E14.5, E16.5 and E18.5. Orange asterisks indicate tissue sections that were excluded from further analysis: a testis at E12.5 and an adrenal gland at E14.5. (**C**) Postnatal stages: P0, P3, P21, P90.

For capture of detailed tissue morphology and alignment of transcriptomic data, we implemented a dual imaging approach. Low resolution images of the H&E-stained sections were captured by the CytAssist instrument (10x Genomics, USA) and were used for spatial alignment of transcriptomic data in SpaceRanger (10x Genomics, USA) (**Fig. S2A, B**). To capture higher resolution images of tissue morphology, we also acquired H&E images at 20X magnification prior to sequencing (**Fig. 1B, C**; *top rows*). Assessment of tissue morphology showed that some of the ovarian tissue was fragmented or folded, a common artifact in fresh frozen tissue sections. However, for all eight timepoints, at least two ovaries were captured with intact tissue morphology. The number of ovaries captured per section differed between the time points, with fetal tissue containing as many as 10 ovaries with good quality morphology. Postnatal timepoints had the most variability in size with 11 ovaries captured at P0 and two complete ovaries captured at P90 (**Fig. 1B, C**; *top rows*). Despite tissue artifacts from sectioning, projection of the integrated cell clusters onto tissue successfully captured highly resolved spatial patterning of cell subtypes both within the ovary and in peripheral tissues (**Fig. 1B, C**; *bottom rows*).

To assess the quality of our sequencing data, we aggregated data from the 2 µm^2^ detection spots on the Visium HD slide into 8 µm^2^ bins and quantified the number of unique molecular identifiers (UMIs) and genes detected in each 8 µm^2^ bin for each timepoint (**Fig. S2 C-J**). Projecting UMI counts onto the tissue showed spatial enrichment of high-quality reads in the ovarian domain relative to surrounding tissue, particularly at late fetal and early perinatal stages (E16.5 to P3; **Fig. S2E-H**). By late postnatal stages (P21 and P90; **Fig. S2I, J**), high UMI counts were concentrated in growing follicles nearest the cortex, which is consistent with increased transcriptional activity associated with primordial follicle activation and subsequent stages of early folliculogenesis. To complement these spatial projections, we also quantified global UMIs and n genes for each timepoint (**Fig. S3K, L**). Across all eight time points, the data show robust transcript capture, with the majority of 8 µm^2^ capture areas containing between 1,000 and 5,000 UMIs. The observed trends in UMI distributions across developmental timepoints were strongly corroborated by the number of genes detected per each 8 µm^2^ spot, which mirrored the UMI count distributions across all eight stages (**Fig. S3K, L)**. Together, the quantitative assessment of UMIs and captured gene distributions along with spatially assessed UMI counts confirmed that our data captured high quality transcripts of the developing mouse ovary. These data further reflect both key biological processes and identifiable technical features, indicating the robustness of our dataset for further investigation.

### Data integration captures spatial and temporal dynamics of developing ovarian cell types

To create a computationally manageable dataset for analysis, we subsampled 50,000 8 µm^2^ bins from each of the eight timepoints, which were combined to create a pool of 400,000 8 µm^2^ bins. Integration of these data using Seurat’s reciprocal PCA method and graph-based clustering identified 34 distinct cell subtypes represented throughout the time-course (**Fig. 2A**). We annotated these clusters using a combination of spatial distribution of known marker genes (**Table 1**) and co-localization of marker gene expression with generated clusters. For example, oocytes were first identified with canonical markers *Ddx4* and *Dazl*. Then meiotic, quiescent primordial and active oocyte clusters were distinguished using spatial gene expression of *Stra8*, which marks entry into meiosis (fetal)(*33*, *43*), *Sohlh1*, a marker of primordial oocytes(*31*), and *Zp3* which becomes expressed in oocytes of primary follicles (postnatal)(*46*). To differentiate somatic cell populations of the ovary, we used canonical markers *Foxl2* and *Runx1* for pre-granulosa cells (PGs), *Up3kb* and *Lhx9* for the ovarian surface epithelium (OSE), and *Col1a1* and *Nr2f2* for stromal cells. PG clusters were further refined using markers of active follicles *Cyp19a1* and *Nr5a2*. Beyond ovary-specific cell types, canonical markers of vascular (*Emcn*, *Pecam1*), immune (*Adgre1*, *Cd86, Cd68*) and lymphatic (*Prox1*, *Lyve1*) cells were used to distinguish between these parallel-patterned clusters. A comprehensive list of the markers used for preliminary identification of cell types is presented in **Table 1**. Of note, we opted to name the clusters based on the ultimate outcome of the cell types. Since all stages were integrated together, our cluster annotations might in some cases be confusing, as spots from the same clusters represent the final differentiated cell type in the older stages, but not their progenitors in the early fetal stages. An example of this is the extraovarian rete cluster (EOR), which at E12.5 and E14.5 captures the Wolffian duct and mesonephric tubules. Because epithelial cells of the mesonephric duct and tubules that remain in the female ultimately form the EOR, we chose to label the cluster “EOR”.

**Figure 2.**
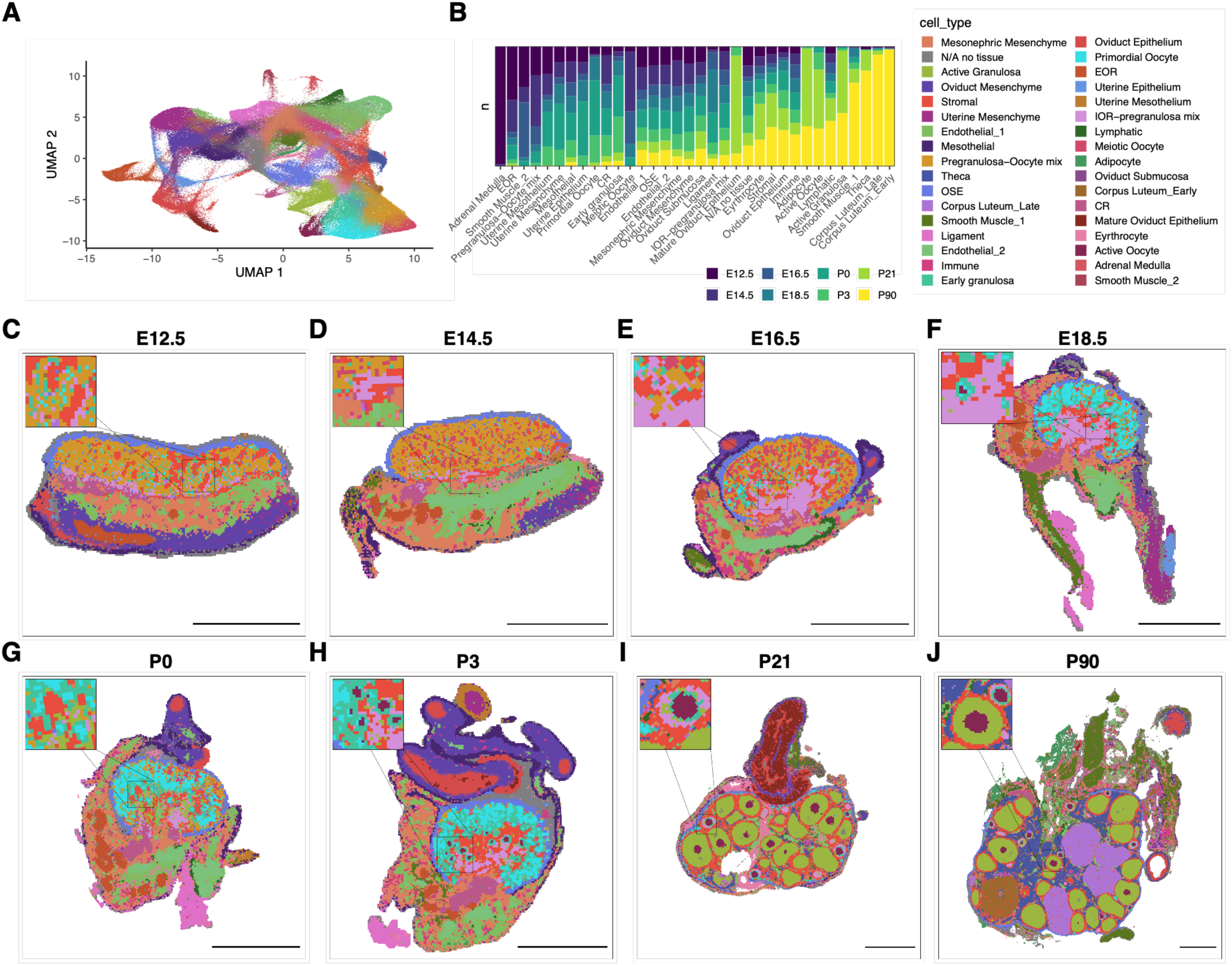
Spatial cluster integration captures dynamic patterning of ovarian cell types across ovarian development. (**A**) UMAP projection of 34 clusters identified using integration of 400,000 subsampled 8 µm^2^ bins across all eight timepoints. *Abbreviations*: OSE, ovarian surface epithelium; IOR, intraovarian rete; CR, connecting rete; EOR, extraovarian rete. (**B**) Bar plot depicting proportion of 8 µm^2^ bins assigned to each cluster at each stage. “Cell_type” displays the 34 annotated clusters used in the UMAP and cluster projections. (**C - J**) Projections of the 34 identified clusters from the integrated dataset onto individual ovaries from each timepoint: (**C**) E12.5, (**D**) E14.5, (**E**) E16.5, (**F**) E18.5, (**G**) P0, (**H**) P3, (**I**) P21, (**J**) P90. The insets in the top left corner show blow-ups of a region within the image, illustrating the high spatial resolution of the clusters. Clipping masks were applied to the background to isolate each ovary from the rest of the image. *Scale bars, 500 μm*.

**Table 1.**
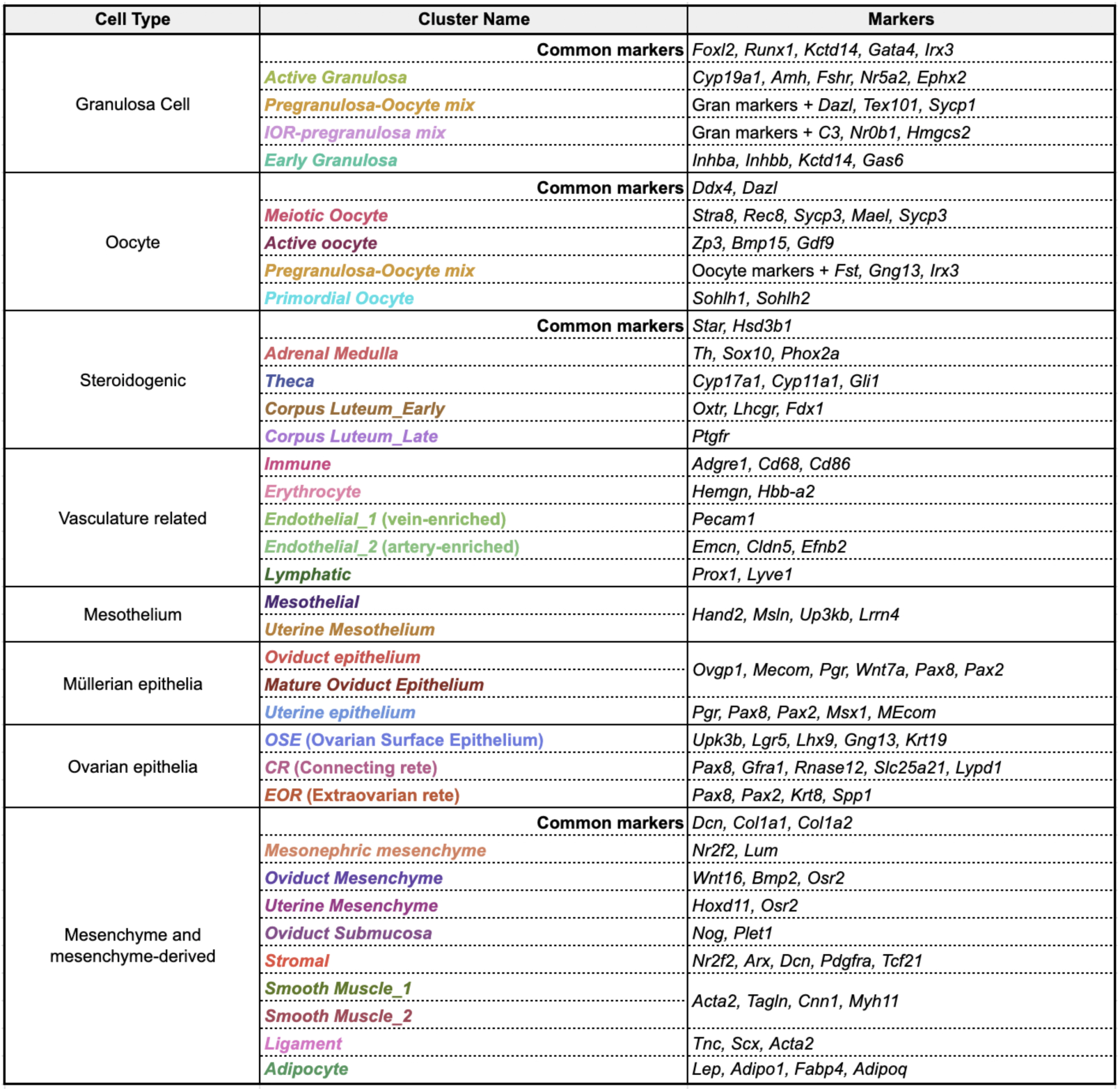
List of cell type markers used to guide cluster annotations.

In addition to spatially informed cluster annotation, the integration of the dataset identified clusters present only at specific windows of ovarian development. For instance, the “adrenal medulla” cluster was found only at E12.5 (large *blue bar* in **Fig. 2B**), consistent with the fact that the adrenal primordium has not yet fully separated from the gonad at this stage (*47*). In contrast, the early and late corpus luteum (CL) clusters were represented only at P90 (*larger yellow bar* in **Fig. 2B).** This is consistent with CLs being a post-ovulatory endocrine structure, and with the first ovulation typically occurring after P21 in CD-1 mice. Of note, we found a small proportion of spots in P21 ovaries assigned to the “corpus luteum late” cluster (∼1% of total spots), which we hypothesize is a sign of follicle atresia (similar to the luteolysis occurring in late CLs), as spatial evaluation showed that these spots were found in smaller, collapsed follicles in these samples.

To assess whether spatial patterning of the clusters was biologically significant, we projected the clusters onto the tissue (**Fig. 2C-J**). Spatial projection of the 34 identified clusters revealed clear, specific, and highly resolved patterning. Clear boundaries between the ovary and peripheral tissue were discernible with the ovarian domain, mesonephros, and oviduct displaying distinct patterning of non-overlapping clusters. Not only did these data discern specific regions and cell types of the ovary based on transcriptomic profiles, but the integrated dataset also captured spatially dynamic transcriptional patterns in the ovary itself across development. At E12.5 and E14.5 the ovarian domain was populated with *early granulosa*, *pre-granulosa/oocyte mix*, and *meiotic oocyte* clusters (**Fig. 2C, D**). By E16.5, the *primordial oocyte* cluster emerged (**Fig. 2E**) as oocytes entered dictyate arrest, and by E18.5 spots assigned to this cluster were densely scattered across the cortex (**Fig. 2F**) highlighting the regionalization into medullary and cortical domains that persisted until P3 (**Fig. 2G, H**). By P21 and P90 most of the previously observed clusters had disappeared and were replaced with clusters associated with the mature ovary, namely: active granulosa cells, active oocytes, theca and corpus luteum (**Fig. 2I, J**).

Together these data indicate that the subsample used for generation of the integrated spatial transcriptomic dataset robustly captured diverse cell subtypes present across ovarian development, that our cluster annotations accurately identify ovarian cell subtypes, and that integration allowed for identification of cell subtypes uniquely represented at specific points of time during ovarian development. This spatial dataset also displayed with unprecedented detail the association between unique gene expression signatures and spatial patterning of ovarian cell types.

### An *in situ* gene expression library at 2 µm resolution

One major advantage of HD spatial transcriptomics is the ability to observe gene expression at near single-cell scale in the context of the tissue. The integrated spatial dataset is in essence a near transcriptome-wide *in situ* gene expression library that one can query for almost every gene expressed in the ovary. While the 10x Visium platform offers a minimal resolution of 2 µm^2^ capture areas – approaching single-cell scale – the inherent sparsity of the spatial transcriptomic data initially necessitated their combination into 8 µm^2^ bins to capture sufficient signal. However, this aggregation came at the cost of spatial resolution. To reclaim high-resolution insights while maintaining data density, we applied Kernel Density Estimation (KDE) using the SAINSC workflow (*48*), which fit a Gaussian kernel to the 2 µm^2^ expression data. This smoothing approach yielded refined 2 µm^2^ expression profiles that offered superior spatial fidelity compared to 8 µm^2^ binning, as demonstrated by the localized expression of *Dazl* in adult oocytes (**Fig. S3A-D).** We subsequently validated that the smoothed 2 µm expression data accurately captured the spatial patterning of known ovarian marker genes at E18.5 (**Fig. 3A)**. Each marker was localized to highly distinct domains where expression was expected: *Up3kb* was localized to the surface of the ovary and surrounding tissue, *Pax8* was localized to the rete ovarii and oviduct epithelia, *Dazl* highlighted the ovarian cortex where most oocytes are localized at E18.5 and *Foxl2* was localized to the ovarian domain (**Fig. 3A**). These are just four selected genes displayed on a single ovary from one timepoint. The total dataset contains eight different time points with – in some cases – nearly a dozen ovaries per section, each of which were sectioned at differing planes, unlocking a wealth of high-confidence gene expression information. Furthermore, the high-resolution data resolved the distinct localization of germ cells (*Dazl*) and PGs (*Foxl2*) (**Fig. 3A**), confirming that the smoothing process preserved critical cellular boundaries.

**Figure 3.**
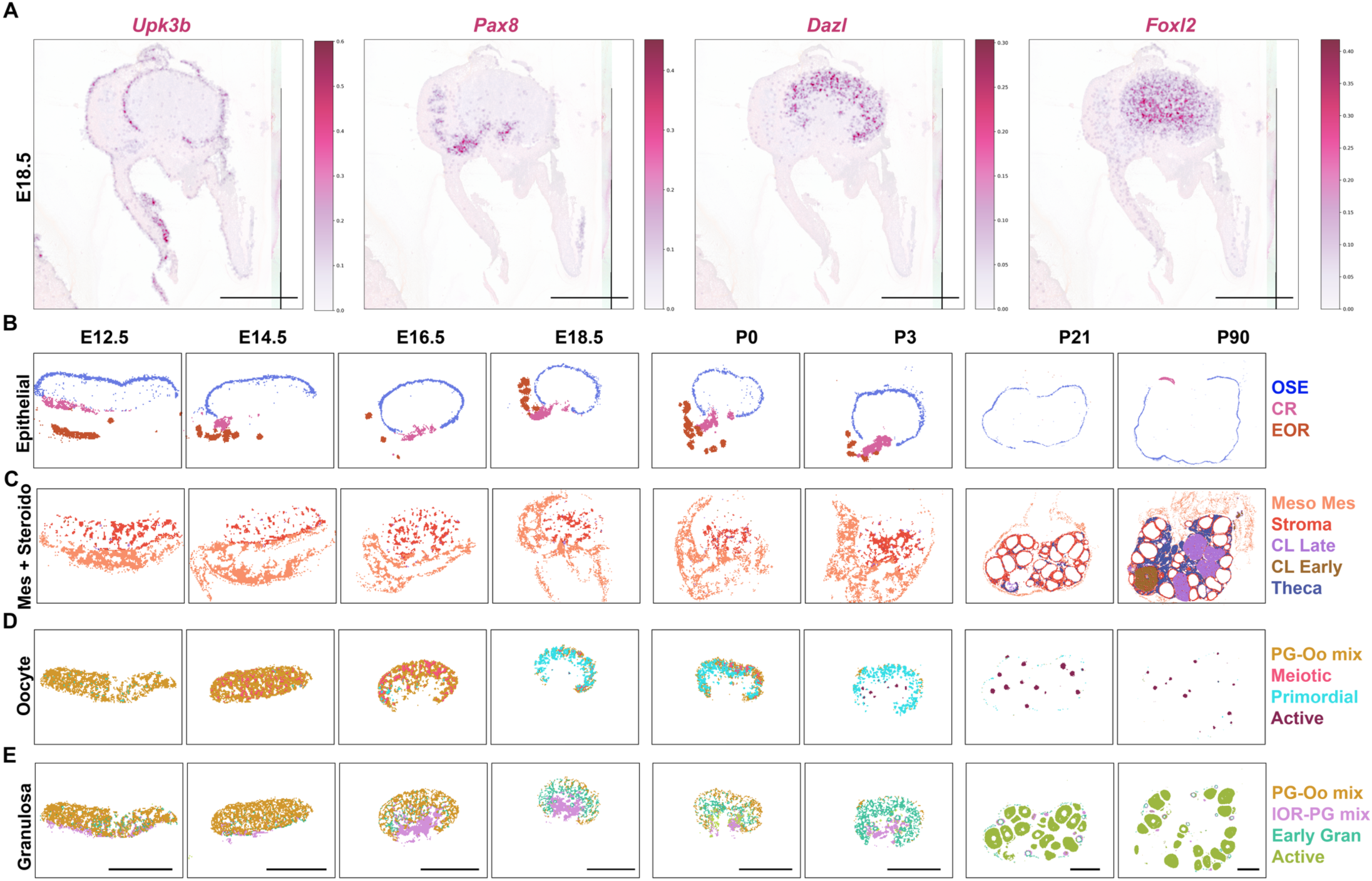
Projection of lineage specific spatial clusters captures elaboration of ovarian components throughout development. (**A**) Smoothed expression of classical ovarian cell types markers at E18.5: *Upk3b* labels the ovarian surface epithelium and the surface of mesonephros-derived tissue; *Pax8* labels the rete ovarii; *Dazl* labels oocytes and *Foxl2* labels pregranulosa cells. (**B-E**) Cluster projection for four distinct lineages across all timepoints; (**B**) Epithelial clusters include ovarian surface epithelium (OSE, *blue*), connecting rete (CR, *rose*), and mesonephric duct (E12.5 only) and extraovarian rete (EOR, all other stages, *terracotta*). CR and EOR were not captured in the representative tissue section at P21, and EOR was not captured at P90. (**C**) Mesenchyme (Mes) and steroidogenic (Steroido) clusters include mesonephric mesenchyme (Meso Mes, *light coral*), which becomes bursa and mesovarium after P21, Stroma (*reddish-orange*), late corpus luteum (CL Late, *mauve*), early corpus luteum (CL Early, *brown*) and Theca (*cobalt*). Theca only arise by P21, surrounding growing follicles, and CLs only arise after ovulation at P90. (**D**) Oocyte clusters include pregranulosa-oocyte mix (PG-Oo mix, *gold*), characterized by expression of both PG and meiotic oocyte marker genes; Meiotic oocytes (*pink*); Primordial oocytes (*cyan*), characterized by markers of meiotic arrest which expand posteriorly from E16.5 to E18.5, and represent a reduced population restricted to the cortex by P21; and Active oocytes (*maroon*), appearing in the center of the ovary by P3, as first wave folliculogenesis begins in the medulla, and occupying a larger medullary compartment ay P21 and P90 with the cyclical waves of adult folliculogenesis. (**E**) Granulosa clusters include pregranulosa-oocyte mix (PG-Oo mix, *gold*), repeated here as it represents a mix of oocytes and PGs; Intraovarian rete-pregranulosa mix (IOR-PG mix, *light purple*), which represents a mix of spots positive for markers of PGs and rete ovarii cells at fetal stages, and populates the most central region of the ovary (ovarian core) from E14.5 to P3, and a subset of granulosa in primary follicles at P21 and P90; Early granulosa (Early Gran, *light teal*), populating the center of the fetal ovary, expanding towards the surface by P3, and forming primordial and primary follicles at P21 and P90; and Active granulosa (Active, *lime green*), which populate the most central region of the ovary at P0, and growing follicles beyond the secondary stage from P3 to P90. Scale bars, 500 μm.

Beyond the advantage this dataset offers as an *in situ* library of ovarian development, we also highlighted how it will enable discovery by performing differential analysis of the top two enriched genes for each cluster using the *FindAllMarkers* function of Seurat (**Fig. S3E**). The resulting heatmap identified cell-type markers not traditionally associated with these annotations, using an unbiased empirical approach. The integrated spatial framework thus enables discovery of novel spatially informed transcriptional signatures across ovarian development. Together, these results demonstrate that our spatial data effectively delineated the complex patterning of diverse ovarian cell types, validating its capacity to resolve the organ’s architecture at both a spatial and transcriptional level.

### Spatial transcriptomics captures the elaboration of discrete ovarian cell populations throughout development and maturation

We next investigated whether the spatial dataset illustrated spatiotemporal dynamics of the developing ovary, by projecting the evolution of clusters associated with structural components of the ovary and discrete cell types, namely: ovarian epithelial structures; mesenchyme, stroma and steroidogenic cells (theca and CLs); oocytes; and granulosa cells. Isolation and projection of the clusters associated with each of these four ovarian components across the eight time points, we were able to more clearly visualize the elaboration of each component from their early distributions into their mature forms (**Fig 3B-E)**.

We first examined the epithelial compartments of the ovary, focusing on the elaboration of the rete ovarii (RO) and the ovarian surface epithelium (OSE). While the RO is retained and significant in the adult ovary (*29*), it is notoriously difficult to access during dissection and cryosectioning due to its location in the perigonadal fat pad (*49*), thus it was not surprising that we captured very little RO at P21 and P90 (**Fig. 3B**). However, we detected two spatial clusters associated with distinct regions of the RO, the extraovarian (EOR) and connecting rete (CR) across all developmental stages up to P3. Remarkably, clusters associated with the RO were present as early as E12.5, indicating that distinct transcriptional landscapes emerge among the different RO components during early development. RO clusters remained transcriptionally distinct throughout development and did not display significant shifts in cluster identity, maintaining their specialized cell type association as the organ matured. However, we could note a shift in the spatial localization of the EOR over time. While initially located near the mesonephros/ovary boundary, by E18.5 the convoluted structure of the EOR emerged and was located medially to the CR, matching previous morphogenic studies of the RO (*29*, *49*). In contrast, the OSE remained localized to the surface of the ovary, with no overt changes in transcriptional representation or cluster identity across development.

We next observed that the clusters representing the mesenchyme / stromal component remained largely stable throughout ovarian development (**Fig. 3C**), exhibiting minimal shifts in cluster identity in the spatial projections over time. The mesonephric mesenchyme cluster was localized to the mesonephros throughout development, and later populated the connective tissue surrounding the ovary, closely associating with the mesovarium and ligaments (**Fig. 3C**). Similarly, the stromal cluster remained regionalized to the internal ovarian domain through much of development (E16.5 – P3). Interestingly, while these stromal progenitors were present throughout ovarian development, they exhibited a distinct shift in late postnatal development from the central ovarian domain to the layers directly adjacent to granulosa cells of growing follicles by P21 and P90, consistent with their role in forming thecal layers during folliculogenesis. The theca cluster itself was not observable until P21, consistent with the onset of steroidogenesis as secondary follicles only emerge after P3. CL clusters were only detectable at P90, consistent with their post-ovulatory nature.

In stark contrast to the mesenchyme, stromal and epithelial components, which maintained stable cluster identities, oocytes and granulosa cells displayed significant shifts in both spatial patterning and cluster identity over time. In oocytes (**Fig. 3D)**, a *meiotic oocyte* cluster emerged during early fetal development between E12.5 and E16.5. – while these cells broadly populated the ovarian domain up to E14.5, by E16.5 this cluster became more restricted to the surface of the ovary. This period also marked a transition in cluster identity from *meiotic oocytes* to arrested *primordial oocytes*, a shift that began in the anterior region and propagated to the posterior pole by E18.5, matching previous findings of an A-P wave of meiosis (*13*, *43*). By P3, the surface of the ovary was exclusively populated by *primordial oocytes*, while *active oocytes* emerged within the central domain in a pattern consistent with the first wave of folliculogenesis occurring in the medulla (*32*). Finally, in late postnatal development (P21–P90), the inner medulla was occupied by *active oocytes,* while *primordial oocytes* remained restricted to the outer periphery at much lower abundances than was observed in earlier stages (**Fig. 3D**). As such, projection of oocyte clusters across development delineated oocyte dynamics that were defined by a coordinated shift in both transcriptional identity and spatial localization, reflecting the divergent fates of germ cells deposited within the medullary and cortical domains of the maturing ovary.

Intriguingly, the shifts in transcriptional identity and spatial localization observed in oocytes were recapitulated by granulosa cells (**Fig. 3E**). As early as E12.5, two transcriptionally distinct populations were already present: *early granulosa* which populated the ovarian domain more generally and *IOR-pregranulosa mix* (named as such due to mixed expression of PG marker *Foxl2* and the intraovarian rete marker *Pax8*) localized primarily at the ovary-mesonephros boundary (**Fig. 3E**). By E16.5, these populations regionalized into two discernable domains, with *early granulosa* localized to the ovarian surface and *IOR-pregranulosa mix* occupying the central ovarian domain, a pattern that continued to resolve through E18.5. At P0, a significant medullary transcriptional shift occurred, resulting in a central domain populated by *both early* and *active granulosa* cells. Interestingly, at the onset of first wave folliculogenesis at P3, it is the *early granulosa* cells, rather than the *active granulosa* population that directly surrounded the growing follicles. Between P3 and P90, we observed a final spatial reorganization, where the *IOR-pregranulosa mix* shifted from a central location at P3 to the periphery of the mature ovary (P21 and P90) and were found in association with primary follicles. This change in spatial distribution was likely not a dispersion of these cell types from the medulla to association with primary follicles; rather, this probably indicated that cells in the *IOR-pregranulosa mix* in the fetal and early perinatal periods shared a transcriptional profile with the active granulosa cells of primary follicles in the mature ovary. Meanwhile, the *active granulosa* cluster previously co-occupying the central domain with *IOR-pregranulosa mix* was more closely associated with follicles that had progressed to secondary or antral stages by P21 and P90 (**Fig. 3E**). Exploration of the spatial and transcriptional dynamics of granulosa cells in the ovary during development thus indicated that granulosa cells were regionalized early in development and that they displayed distinct region-specific transcriptional landscapes.

### Differential gene expression reveals early expression of follicle activation genes in center PGs

Following validation that our dataset captured the spatial and transcriptional diversity of the ovary across development, we next examined patterning events that inform the establishment of adult ovarian function, particularly those governing the establishment of the ovarian reserve. The regionalization of the ovary into cortical and medullary domains is a critical feature of ovary development, which results shortly after birth in the establishment of the quiescent ovarian reserve in the surface (cortex), and first wave folliculogenesis in the center (medulla) (*24*, *25*, *32*). Here we used our spatial transcriptome dataset to begin to identify spatial and molecular components that underly this regionalization.

While oocyte clusters did not display discrete transcriptional signatures along the cortical-medullary axis, we identified two transcriptionally and spatially distinct PG clusters in the fetal ovary that coincided with the emerging central and surface domains. These two populations were already transcriptionally distinct by E16.5 (*light purple* cluster in **Fig. 3E**). Whole-mount immunofluorescent (IF) labeling of PGs with RUNX1 and germ cells with DDX4 revealed distinct morphological domains in the ovary at E16.5. While the surface was populated by more sparsely arranged PGs interspersed with dense collections of oocyte nests (**Fig. 4A***, inset 1*), PGs in the center were arranged in a dense scaffold (**Fig. 4A**, *inset 2*). These center PGs displayed higher intensity RUNX1+ signal and appeared to anchor the ovigerous cords extending toward the surface of the ovary at E16.5.

**Figure 4.**
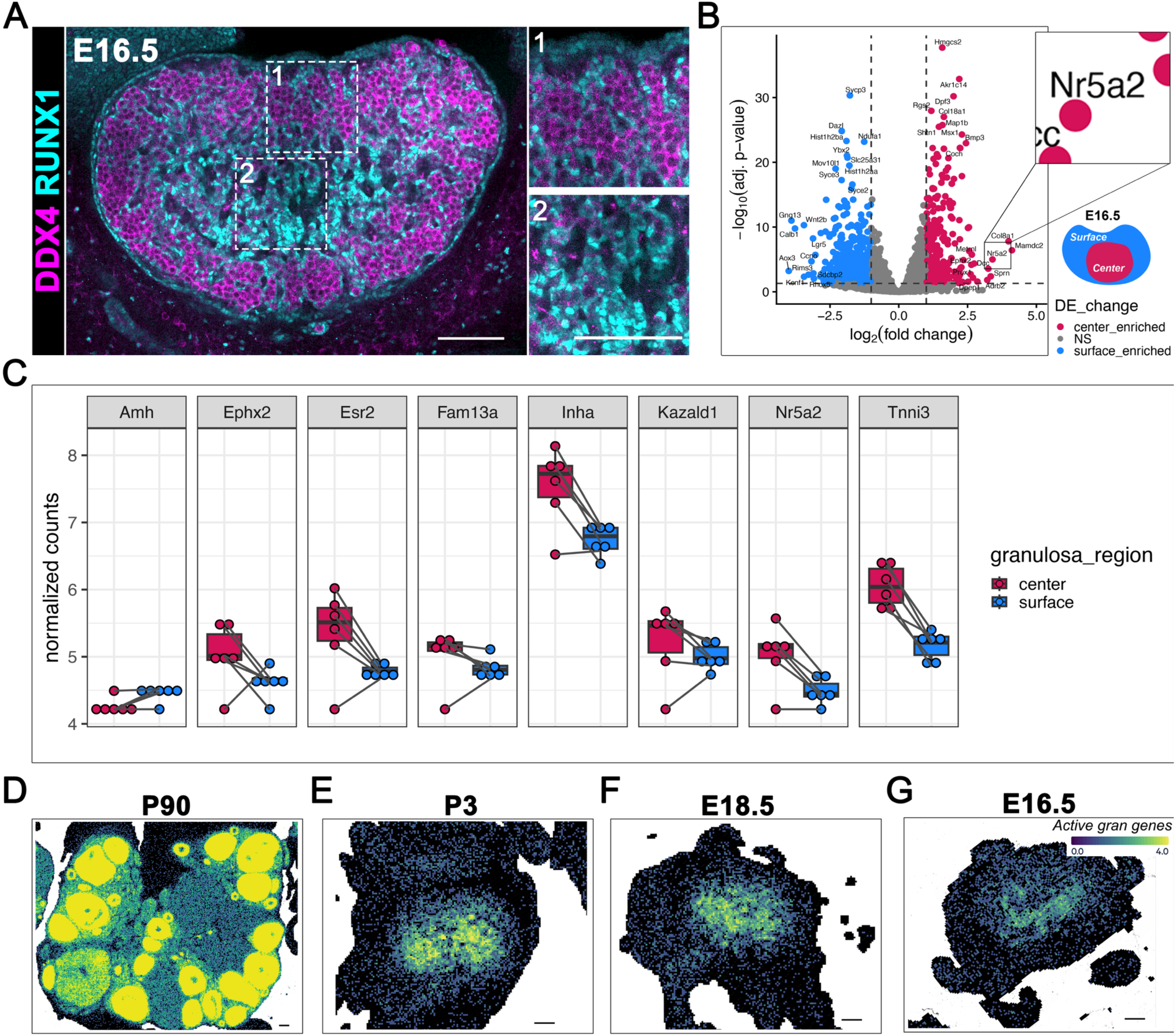
DGE reveals activation-poised PGs in the central domain of the ovary by E16.5. (**A**) Optical section from a representative confocal Z-stacks of a whole-mount E16.5 ovary immunostained with DDX4 (*magenta*) and RUNX1 (*cyan*). Inset 1 shows the surface of the ovary, inset 2 shows the center. Scale bar, 100 µm. (**B**) Volcano plot of differential gene expression between center enriched (*active granulosa*, and *IOR-pregranulosa mixed* clusters) and surface enriched (*oocyte-granulosa mix* and *early granulosa* clusters) 8 µm^2^ bins. Bins were deconvolved for ≥80% granulosa identity and pseudobulked prior to analysis. Inset highlights *Nr5a2* (top right). Bottom right illustrates center and surface regions of the E16.5 ovary. (**C**) Box plot of normalized counts of genes from active signature between the center and surface at E16.5: *Amh, Ephx2, Esr2, Fam13a, Inha, Kazald1, Nr5a2,* and *Tnni3*. Dots represent replicate ovaries used for analysis, paired dots represent surface and center sampled from the same replicate. (**D–G**) Compiled active signature genes (sum *Amh, Ephx2, Esr2, Fam13a, Inha, Kazald1, Nr5a2,* and *Tnni3* expression) projected onto P90 (**D**), P3 (**E**), E18.5 (**F**) and E16.5 (**E**) ovaries. Scale bar, 100 µm.

To discover major drivers of the transcriptional differences between surface and center PGs in the fetal ovary, we performed differential gene expression (DGE) analysis across three timepoints: E16.5, when spatial differences emerge; at E18.5, when regionalization is established; and at P3 when medullary follicles activate and undergo first wave folliculogenesis. Embedding of many ovaries for each timepoint allowed us to aggregate spots from each ovary section as a pseudobulk replicate (**Fig. S4A**) to perform high-confidence DGE analysis. Ovaries sectioned at an ideal plane through the central and surface domains were selected for each timepoint and treated as pseudobulking replicates (E16.5, N = 6; E18.5, N = 6; and P3, N = 7). A technical limitation of this approach arose from the stochastic overlap of the capture area with ovarian tissue and the size of the bins themselves. This resulted in some 8 µm^2^ bins containing mixed cell types (**Fig. S4B-D**). To overcome this limitation, we performed spot deconvolution using robust decomposition of cell type mixtures (RCTD). We used snRNA-seq data that we obtained from E16.5 and P0 ovary/rete ovarii complexes as a reference dataset (**Fig. S5**). This strategy allowed us to select bins enriched for RNA from PGs by estimating the proportion of RNA contributed from different cell types in each 8 µm^2^ bin. Deconvolution was conducted independently for each spatial dataset (E16.5, E18.5, and P3) and RCTD was run in doublet mode, reflecting the expectation that individual bins would span no more than two cell types.

For each bin, RCTD estimated the relative RNA contribution of reference-defined cell types. We retained only bins predicted to contain ≥80% RNA from granulosa lineage populations, thereby minimizing contamination from non-granulosa cell types (**Fig. S4B-D**, compare *rctd_filtered=FALSE* to *rctd_filtered=TRUE*). Granulosa bins were further stratified based on spatial annotation: bins classified as *active granulosa* or *IOR–pregranulosa mix* were designated as central, whereas bins annotated as *early granulosa* or *oocyte–granulosa mix* were designated as surface. Differential expression analysis was then performed using a pseudobulk framework in which center- and surface-classified bins were aggregated within each ovary to generate biological replicates. This approach enabled DGE comparisons with 6–7 replicates per stage, a level of replication rarely achievable in single-cell–scale studies. Differential expression was determined using DESeq2 at E16.5 (**Fig. 4B**), E18.5 and P3 (**Fig. S6**) (*50*).

Among the genes enriched within center PGs at E16.5, *Nr5a2* emerged as a prominent differentially expressed transcription factor (**Fig. 4B**). *Nr5a2* is a nuclear receptor used as a canonical marker of granulosa cell differentiation, a process typically associated with follicle activation. Its enrichment within centrally located PGs several days prior to the onset of postnatal folliculogenesis suggests that these cells acquire transcriptional competence well before morphological activation. To determine whether this pattern extended beyond *Nr5a2*, we examined additional genes previously associated with granulosa cell (GC) activation. Previous work identified a group of nine genes including *Nr5a2* that underly GC activation: *Slc18a2*, *Amh, Ephx2*, *Esr2*, *Fam13a, Inha*, *Kazald1*, and *Tnni3* (*37*, *51*). Collectively, these genes displayed elevated expression within the central domain relative to the ovarian surface (**Fig. 4C**; **Fig. S6**), revealing a spatially restricted transcriptional program consistent with early developmental priming. To validate the regional specificity of this activation-associated program, we projected the aggregated gene set onto our spatial transcriptomic data. Individual patterns of expression of these genes are displayed in **Fig. S7**. By P90, robust enrichment was observed within growing follicles across multiple stages of follicular development, confirming that this program characterizes functionally active granulosa cells (**Fig. 4D**). A similar spatial pattern emerged at P3, where enrichment localized to the central domain, coinciding with granulosa differentiation during first-wave folliculogenesis (**Fig. 4E**). Unexpectedly, this central transcriptional enrichment was already evident in the fetal ovary at both E18.5 (**Fig. 4F**) and E16.5 (**Fig. 4G**), well before the onset of follicle formation. The presence of this activation signature within centrally located PGs during fetal life suggests that these PGs acquired competence for activation prior to morphological differentiation. This intriguing finding indicates that spatial patterning of PGs in the center of the ovary may anticipate future follicle activation.

### Center PGs express markers of follicle activation before follicle assembly

Having identified in our spatial transcriptome a subdomain within the center of the ovary that expressed markers of follicle activation in the fetal ovary at E16.5, we next sought to validate this finding using orthogonal methods, including IF for protein detection and RNAscope (ACDBio, Inc., USA) for *in situ* detection of RNA. We first examined expression of the canonical granulosa activation marker *Nr5a2*, as it was one of the top enriched genes in center PGs at E16.5 (**Fig. 4B**). As expected, our spatial dataset confirmed detection of *Nr5a2* in active granulosa cells of growing medullary follicles at P3, when FWA was well under way (**Fig 5A**). Examination at earlier stages revealed that *Nr5a2* was already detected at E18.5, specifically in the center of the ovary. Notably, this expression domain was also observable at E16.5, coinciding with the regional compartmentalization of the PG spatial clusters (**Fig. 3E, E16.5; Fig. 5A**).

**Figure 5.**
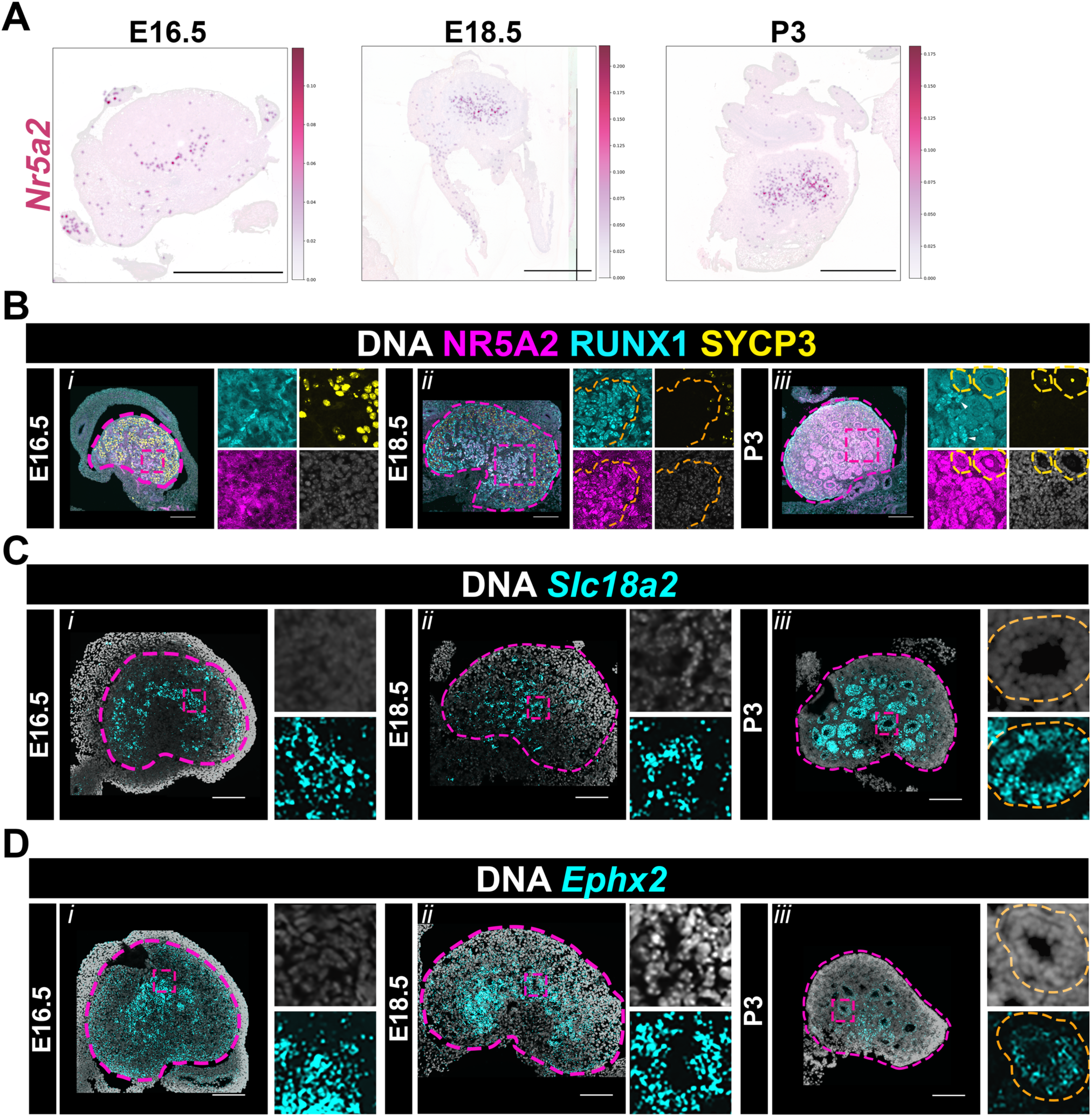
Genes of the active signature are captured with spatial transcriptomics, IF and RNAscope in the central domain of the fetal and perinatal ovary. (**A**) Smoothed 2 µm expression of *Nr5a2* projected onto E16.5, E18.5 and P3 ovary. Scale bar, 500 µm (**B**) Maximum intensity projection (MIP) from confocal Z-stacks of ovaries collected at E16.5 (*i*), E18.5 (*ii*) and P3 (*iii*), cryosectioned at 10 µm and immunostained for NR5A2 (*magenta*) and RUNX1 (*cyan*), SYCP3 (*yellow*) and counterstained with Hoechst nuclear dye (DNA, *grayscale*). Ovaries outlined in magenta. Core PGs outlined in orange. Follicles outlined in yellow. Insets show split channels of central domain in each image. Scale bar, 100 µm. White arrowheads at P3 indicate NR5A2+/RUNX1+ cells not incorporated into follicles. (**C-D)** Maximum intensity projection (MIP) from confocal Z-stacks of ovaries collected at E16.5 (*i*), E18.5 (*ii*) and P3 (*iii*), cryosectioned at 10 µm for *in situ* hybridization using RNAscope probes for *Slc18a2* (**C**) or *Ephx2* (**D**) (*cyan*) and counterstained with DAPI (DNA, *grayscale*). Outlines: ovary (*magenta*) and growing follicles (*orange*). Insets show split channel enlargements of the center. Scale bar, 100 µm.

We next investigated protein expression using antibodies against NR5A2, which were first validated by specific expression in growing follicles of the adult ovary (**Fig. S8A**). This experiment also confirmed that NR5A2 was not expressed in quiescent PGs of primordial follicles (**Fig. S8A***, yellow outlines*). We next examined NR5A2 expression at developmental timepoints. At P3, IF showed expression of NR5A2 in the central domain, in active granulosa cells where it co-localized with RUNX1 (**Fig 5B***iii*), recapitulating the expected localization of NR5A2 in growing follicles. Consistent with the expression pattern identified in our spatial dataset, we found that NR5A2 protein was also expressed in the fetal ovary, showing strong expression at E18.5 localized to RUNX1+ cells in the center of the ovary (**Fig. 5B***ii*). While expression was lower, we also detected NR5A2 in center PGs at E16.5 (**Fig. 4B***i*). These data indicated that PGs in the center express NR5A2 even before assembling into follicles. Intriguingly, RUNX1+/NR5A2+ PGs in the fetal ovary formed a dense population of PGs that branched radially from the core of the ovary to the rest of the medulla (**Fig. 5B***ii*), anchoring ovigerous cords. This core region was also sparsely populated with germ cells (SYCP3+, *yellow*) compared to the rest of the ovary (**Fig. 5B***ii*). Additionally, we also observed RUNX1+/NR5A2+ cells at P3 that were not incorporated into growing follicles or even near SYCP3+ germ cells (**Fig. 6B***iii*, *white arrowheads*).

**Figure 6.**
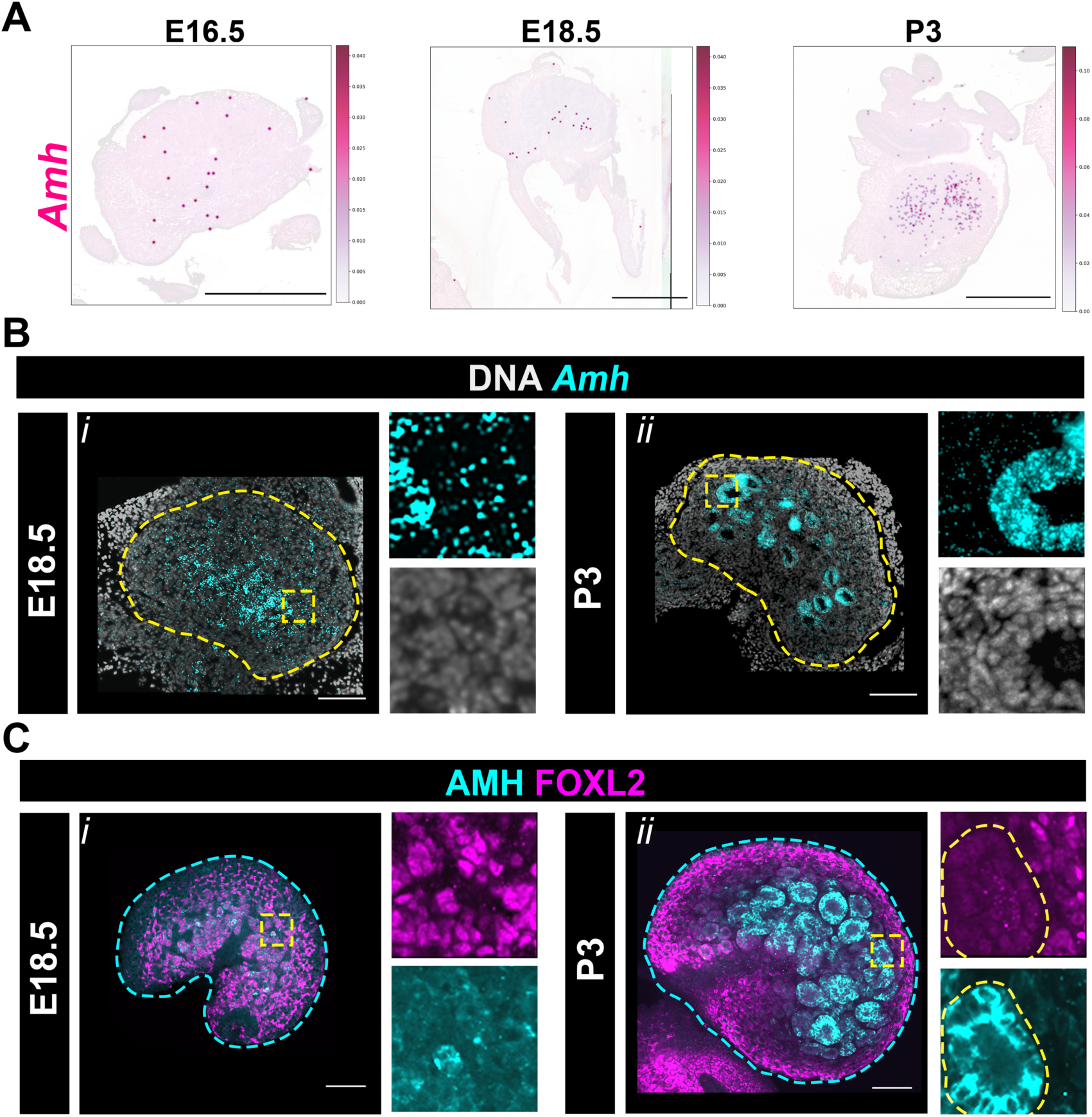
Unexpected expression of anti Müllerian hormone inf the fetal ovary. (**A**) Smoothed 2 µm expression of *Amh* projected onto a E16.5, E18.5 and P3 ovary. Scale bar, 500 µm. (**B**) Maximum intensity projection (MIP) from confocal Z-stacks of ovaries collected at E18.5 and P3, cryosectioned at 10 µm for *in situ* hybridization using a probe for *Amh (cyan*) and counterstained with DAPI (DNA, *grayscale*). Outlines: ovary (*yellow*), insets (*yellow box*) (**C**) Optical section from confocal Z-stacks of a whole-mount E18.5 and P3 ovaries immunostained with FOXL2 (*magenta*) and AMH (*cyan*). Insets of medullary regions (*yellow box*). Outlines: ovary (*cyan*) and growing follicles (*yellow*) Scale bar, 100 µm. (**C**) Maximum intensity projection (MIP) from confocal Z-stacks of ovaries collected at E18.5 and P3, cryosectioned at 10 µm for *in situ* hybridization using a probe for *Amh (cyan*) and counterstained with DAPI (DNA, *grayscale*). Outlines: ovary (*yellow*), insets are enlargements of central region (*yellow box*). Scale bar, 100 µm.

After validating the presence of a known marker of granulosa activation in the adult, perinatal and fetal ovary, we next examined the expression of two other active signature genes using RNAscope. *Slc18a2* was recently identified as one of the first genes to turn on in activating granulosa cells of the perinatal ovary (*37*). However, we were unable to query *Slc18a2* in our spatial dataset and DGE analysis, as it is not included in the default Visium HD probe set. RNAscope probes directed against *Slc18a2* in the adult ovary confirmed robust expression in granulosa cells of growing follicles beyond the primary stage (**Fig. S8B)**. Similar to NR5A2, we found that *Slc18a2* transcripts were enriched in the core domain of the fetal ovary at E16.5 (**Fig. 5C***i*), and that this domain was maintained at E18.5 (**Fig. 5C***ii*). As expected, *Slc18a2* expression was highly enriched in active granulosa cells of medullary growing follicles at P3 (**Fig. 5C***iii*). Importantly, throughout these stages, *Slc18a2* was not detected in the surface of the ovary, where PGs are associated with quiescent oocytes. Thus, the expression of *Slc18a2* transcripts matched the localization of center PG and active granulosa clusters in the spatial dataset. We also investigated the expression of *Ephx2*, another gene that was revealed in our DGE analysis as center-enriched at E16.5, and whose expression was previously mapped to active granulosa cells (*52*). Here again, RNAscope at P3 confirmed localization of *Ephx2* to growing follicles of the medulla (**Fig. 5D***iii*). In the fetal ovary, *Ephx2* was also enriched in the core central domain at E18.5 (**Fig. 5D***ii*) and as early as E16.5 (**Fig. 5D***i*), though we also detected low levels of expression near the surface (**Fig. 5D**). To complete these observations, we also examined the expression profile of five other genes of the active signature (*Inha; Esr2; Tnni3; Fam13a; Kazald1*) (*37*, *51*) in our spatial dataset, and all showed detectable expression in the center of the ovary by E16.5 (**Fig. S7**). Together these data suggest that a small subset of medullary PGs located specifically in the innermost core of the ovary which we will refer to as “core PGs”, are poised for activation long before first wave folliculogenesis or even follicle assembly.

### Unexpected expression of anti-Müllerian hormone in the fetal mouse ovary

Anti-Müllerian hormone (AMH) is the major canonical marker of growing follicles, where it plays a significant role in promoting quiescence of surrounding primordial follicles (*8*, *53*–*55*). AMH is also critical during development for the regression of the Müllerian duct in male embryos, thus it is typically assumed to be absent from fetal ovaries (*56*–*60*). However, given the widespread expression of other markers of granulosa activation in core PGs, we chose to examine the expression of Amh in the perinatal and developing ovary. We first queried our spatial dataset and found that *Amh* was barely detectable until P3, where it was enriched in the medulla, coinciding with growing follicles of the first wave (**Fig. 6A**).

We used RNAscope to examine *Amh* expression further. As expected, RNAscope on adult ovaries showed that *Amh* was specifically enriched in granulosa cells of growing follicles but was absent from quiescent PGs of primordial follicles (**Fig. S8D**) This pattern was recapitulated with IF to detect AMH protein (**Fig. S8A**). Having confirmed that our methods captured known patterning of AMH in the adult, we next validated its expression at P3. RNAscope showed *Amh* transcript localization to growing follicles present in the central region of the ovary (**Fig. 6B***ii*). Whole mount IF at P3 also recapitulated this regionalization, where robust AMH protein expression was detected in active FOXL2+ granulosa cells of growing medullary follicles (**Fig. 6C***ii*). Having recapitulated expected expression of AMH with both RNAscope and IF in adult and perinatal ovaries, we next examined AMH expression in the fetal ovary. RNAscope revealed significant *Amh* expression specifically in the core domain of the ovary at E18.5 (**Fig. 6B***i*), which appeared to slightly extend toward the surface of the ovary. In addition, whole-mount IF also revealed expression of AMH protein in FOXL2+ PGs of the core medullary domain in the E18.5 ovary (**Fig. 6C***i*). Together these data reveal unexpected expression of Amh at both the transcript and protein level in activation-poised core PGs of the fetal ovary by E18.5.

## Discussion

In the present study, we utilized 10x Genomics Visium HD technology (*61*) (10x Genomics, Inc) to construct a single-cell scale spatial transcriptomic time course of the developing mouse ovary across eight critical fetal and postnatal stages. This high-resolution approach enabled the direct integration of transcriptional signatures with precise anatomical regionalization, providing a comprehensive library of *in situ* gene expression. Our dataset captured all principal ovarian cell types in their native context, with biological replication across multiple planes and orientations to ensure high-confidence analysis of spatially distributed cell subtypes. This resource has refined our understanding of ovarian architecture by revealing a previously uncharacterized medullary core domain that emerges by E16.5. Pre-granulosa cells (PGs) within this core express a molecular signature typically restricted to postnatally activated granulosa cells, suggesting they are poised for activation while still in the fetal environment. These findings challenge existing paradigms of ovarian follicle wave specification and establish a new structural framework for investigating the developmental determinants of ovarian health.

### Advancing spatial resolution to define ovarian regionalization

Spatial transcriptomic technologies have enabled the discovery of molecular determinants of tissue architecture across the adult female reproductive system, including the cycling mouse uterus (*62*), the fetal ovary in non-human primates (*63*) and the adult ovary in mice (*64*, *65*), and humans (*66*, *67*). However, the developing mouse ovary has been more challenging to investigate using spatial transcriptomics, as previous investigations were constrained by 55 µm capture spots, which aggregate signals from multiple cell types and preclude precise interrogation of ovarian lineages in the densely packed fetal ovary (*68*).The transition to the Visium HD platform represents a significant technical advancement for resolving complex cellular architecture. By leveraging the near-single-cell resolution of the HD platform, we overcome these limitations, facilitating the definitive mapping of transcriptional signatures to anatomical domains. While the primary focus of this study is the ovary, our dataset includes the mesonephric and Müllerian systems, enabling precise analysis of the spatial molecular determinants of the female reproductive tract across development. To address the technical challenge of transcript capture sparsity at this scale, we implemented the SAINSC workflow to generate smoothed expression profiles, enhancing the visualization of low-abundance transcripts (*48*). Furthermore, we utilized spot deconvolution informed by our reference single-nucleus RNAseq dataset to assign high-quality spatial bins to specific cell types, enabling robust differential gene expression analysis between regional subpopulations. The importance of this spatial framework is evident when contrasted with existing dissociated single-cell datasets (*25*, *26*, *34*, *37*, *41*). In traditional scRNA-seq, PGs were clustered based on global molecular similarities, which likely masked low-level expression of follicle activation-associated genes. In our spatial model, cells clustered based on anatomical location, effectively amplifying the signal of the activation-poised population and revealing molecular regionalization that was previously lost to the averaging effects of molecular clustering in non-spatial datasets. This could explain why the early expression of the follicle activation signature was not identified in fetal scRNA-seq datasets.

### Spatial heterogeneity and lineage specification during ovarian morphogenesis

Ovary morphogenesis is characterized by dramatic structural rearrangements that occur concomitantly with the molecular differentiation of somatic and germ cell lineages (*1*, *10*, *24*, *29*, *32*). We previously identified ovary folding as a key hallmark of this process, wherein the elongated fetal gonad transitions into a crescent shape (*29*). Our current spatial transcriptomic data confirm that this folding between E14.5 and E16.5 is tightly correlated with the regionalization of the ovary into medullary and cortical compartments. This compartmentalization is underpinned by the specification of distinct PG populations that originate from different sources and at different times. The PGs originating from the coelomic epithelium (CE) prior to gonadal sex determination are the first to associate with germ cell cysts, and settle predominantly in the medulla (*24*, *25*, *41*). A second population of PGs ingresses from *Lgr5*+ progenitors of the ovarian surface epithelium from E14.5 onward, contributing to the cortical primordial follicles that constitute the lifelong ovarian reserve (*15*, *24*, *27*, *69*). While these two “waves” arise from the epithelium at the ventral surface of the ovary, a recently discovered third PG lineage derives from supporting-like-cells (SLCs) on the dorsal side (*70*). Specified as early as E10.5, SLCs contribute to the rete ovarii and to medullary PGs, representing a transient population active during early folliculogenesis (*35*). Our study adds a new layer of complexity to this radial architecture by revealing further sub-compartmentalization within the medulla. We identified a core population of PGs in the ovarian center that express genes associated with follicle activation as early as E16.5. Although initial clustering grouped these core PGs with *Pax8*+ cells of the intraovarian rete (IOR) due to their proximity, spatial deconvolution revealed no overlap between the "activation-poised" transcriptional signature and the SLC marker *Pax8*. This suggests that core PGs are molecularly distinct from SLCs, though it does not preclude an SLC origin. Ultimately, lineage tracing and high-resolution trajectory analysis will be required to determine whether core PGs descend from the CE or the SLC lineage.

### The activation-poised medullary core

One of the most significant findings of this study is the identification of a third biological compartment within the ovary: the central medullary core. Spot deconvolution followed by pseudobulk differential gene expression analysis revealed that PGs within this domain express genes of the "activating follicle signature" as early as E16.5. This signature includes several genes identified in two previous studies as markers of perinatally activating granulosa cells: *Nr5a2*; *Slc18a2*, *Tnni3*, *Inha*, *Esr2*, *Fam13a*, *Amh*, and *Ephx2* (*37*, *51*). Intriguingly, these studies did not detect robust expression of this gene expression signature until after birth. Both studies primarily used the C57Bl/6 mouse strain, while our study utilized the outbred CD1 strain, and previous research has demonstrated that the timing of follicle assembly and activation can vary significantly between strains. Thus temporal differences may be attributable to mouse strain differences (*71*). In addition, the spatial nature of our dataset likely allowed us to capture the early expression of the active signature more effectively than dissociated scRNA-seq, as the concentration of these cells in a dense medullary core prevented the signal from being diluted or lost during clustering. We validated the expression of several key components of the activation signature using orthogonal methods.

Immunofluorescence confirmed the presence of NR5A2 protein in the central region as early as E16.5, while RNAscope localized *Amh*, *Slc18a2* and *Ephx2* transcripts to the same domain. Intriguingly, RUNX1+/NR5A2+ core PGs adopted a distinct morphology from the rest of the medullary PGs, forming dense cords of cells that did not incorporate into follicles even after the onset of first wave folliculogenesis at P3. Further studies are required to precisely assess the nature of these cells and whether they are in fact PGs that are eventually recruited to growing follicles. Core PGs may remain extrinsic to follicles, perhaps giving rise to the peripubertal androgen-producing signatures recently reported in the lineage trace and scRNA-seq examination by Yin and Spradling (*26*). Another possibility is that core PGs contribute to the subpopulation of follicles recently identified by Soygur *et al*. that activate during the first wave of folliculogenesis but persist throughout the reproductive lifespan (*33*). From a translational perspective, it will be interesting in the future to integrate these data with recent spatial transcriptomics of non-human primate fetal ovaries (*63*) to determine whether the core medullary domain, core PGs, and their potential contribution to female fertility and reproductive health are conserved.

### Spatial Determinants of Medullary Specification

A central question arising from our observations is how spatial localization is translated into molecular signaling to define regional cell subtypes within the ovary. Activation-poised core PGs (within the IOR-PG cluster) were revealed as an NR5A2+ / RUNX1+ subset of the medullary lineage, yet they exhibit a transcriptomic profile distinct from their medullary counterparts (clustered with the IOR-PG mix). Several factors may define their differential expression. The earliest specified CE and SLC-derived PGs settle deepest in the medulla and are the first to be exposed to the systemic networks arriving through the hilum (*24*, *35*, *42*, *70*, *72*), suggesting the dorsal niche of the fetal ovary may promote the activation-poised signature. The medulla is characterized by high vascular density (*10*, *29*, *73*) and early innervation (*74*, *75*), providing a conduit for systemic growth factors and hormones that are less accessible to the cortical compartment. The physical folding of the ovary, which internalizes the mesovarium and its associated tissues (26), may create a unique microenvironment where mechanical tension and biochemical signaling converge. While the cortical ECM is known to promote quiescence in the adult ovary, the local ECM deposition in the medullary core during fetal development may provide inductive cues that support the activation-poised core PGs. One of the top enriched genes in core PGs was *Col8a1*, a short-chain non-fibrillar collagen known for its role in regulating tissue stiffness and vascular smooth muscle cell proliferation and migration (*76*). Future research is necessary to uncover how ECM regulation impacts formation of the medullary niche in the fetal ovary. Additionally, the rete ovarii is intimately associated with the medullary core. The rete ovarii has been implicated in meiotic progression and follicle formation, and it is possible that paracrine factors from this epithelial structure contribute to the priming of core PGs toward activation (*49*, *77*). Future work integrating this spatial dataset with proteomic approaches and recent cis-regulatory profiling studies (*39*, *78*) will be essential for defining the gene regulatory networks controlling regional cell subtype specification in the developing ovary.

### Significance of ovarian regionalization for establishment of the ovarian reserve

Follicles assembled in the medulla during fetal life activate shortly after birth and are mostly depleted by puberty (*31*, *33*). The existence of this activation-poised population that does not directly contribute to offspring suggests a broader physiological or structural purpose. Core PGs may serve as a structural organizing component for the radial architecture of the ovary, providing a scaffold upon which the rest of the ovarian tissue is patterned, and facilitating the inward folding and subsequent regionalization that characterizes morphogenesis (*29*, *32*). Core PGs could also contribute to the onset and/or propagation of the signals that trigger first wave folliculogenesis. Understanding how core PGs emerge and why they become poised for activation will provide critical insight into the establishment and maintenance of the reserve during the perinatal period. A growing theory is that medullary follicles may act as a shield for the cortical reserve (*79*). By responding early to external perturbations and systemic signals, these follicles may absorb potential insults, thereby protecting the finite pool of quiescent follicles at the surface. Among genes of the follicle activation signature, the detection of AMH in core PGs at E18.5 was particularly remarkable. While AMH is best known for its essential role in controlling Müllerian duct regression during male development (*56*, *58*, *59*), it is also a canonical marker of ovarian follicle activation in perinatal and adult females (*33*, *53*, *54*, *80*). However, AMH is typically assumed to be absent in the fetal ovary, and to remain undetectable until medullary follicles begin to grow postnatally (*60*). AMH is a potent inhibitor of follicle activation in the adult ovary, considered the main gatekeeper of the ovarian reserve (*51*, *81*). AMH produced by core PGs may reinforce the quiescence of cortical primordial follicles right before birth to prevent their recruitment during first wave activation. Future studies will determine the activity of AMH produced by core PGs of the fetal ovary and whether this plays an important role in specification of regional subtypes, perinatal follicle activation, and establishment of the ovarian reserve. Defects in this shielding mechanism could have lasting effects on reproductive longevity. For example, if the medullary core fails to establish this protective boundary or if the ’poised’ state is disrupted, cortical PGs may be exposed to inappropriate activation signals, depleting the primordial reserve before reproductive life even begins. This provides a potential developmental mechanism for cases of primary ovarian insufficiency with unknown etiology (*3*, *82*–*84*), linking fetal spatial patterning directly to adult ovarian health.

While we have focused on the medullary core subpopulation, the potential of this dataset for uncovering new spatial actors across all cell types and developmental stages is vastFuture work should address the specific mitogenic signals that trigger the transition of poised PGs to proliferation and the roles of surrounding cells and tissues in regulating these events. By establishing a comprehensive blueprint of the gene regulatory networks controlling regional cell subtype specification, we can move closer to unlocking the etiology of POI and identifying therapeutic targets to preserve female fertility and extend reproductive longevity. The synthesis of spatial patterning and molecular signatures provided here is a critical first step toward a holistic understanding of the mammalian ovary as a highly dynamic organ.

## Materials and Methods

### Mice

Outbred CD-1 mice were obtained from Charles River, Inc. (USA) and bred in our colony to produce all timepoints in this dataset. To collect ovarian tissue at specific developmental stages, triad timed matings were set up using a single male and two two-month-old virgin females. Timed-mated dams were checked every morning for vaginal plugs. The date of vaginal plug was treated as embryonic (E) day 0.5. Mice were housed in an AALAC-accredited barrier facility at the University of Colorado Anschutz Medical campus under a 14-hour light:10-hour dark cycle, at 22.2°C +/- 1.1°C with 40% +/- 10% humidity. Mice were fed with a standard ad libitum diet (2920X, Teklad, Inc. USA) and watered with hyperchlorinated water (2 - 5 ppm). All mice were handled and cared for following accepted National Institute of Health guidelines. All experiments were conducted with the approval of the University of Colorado Anschutz Institutional Animal Care and Use Committee (IACUC protocol # 01262).

### Fetal tissue collection

All tools were disinfected with 70% ethanol and sprayed with RNaseZap (Invitrogen, USA) before use. 1X phosphate buffered saline without calcium or magnesium (PBS -/-) was made using H_2_O with diethyl pyrocarbonate (DEPC-H_2_O) to prevent RNase activity during sample procurement. At the desired gestational age, pregnant dams were euthanized by the isoflurane drop method followed by cervical dislocation per university approved IACUC protocol #01262. Embryos were harvested into 1X DEPC-PBS -/-on ice. Both ovary / mesonephros complexes from each female embryo were quickly dissected and added to a droplet of optimal cutting temperature compound (OCT) on a petri dish. This allowed removal of excess DEPC-PBS before transferring ovary-mesonephros complexes to a cryomold. Our goal was to place as many ovary/mesonephros complexes as possible into one cryomold (**Fig. 1**). Thus, two pregnant dams were used for E12.5, E16.5, and E18.5, and three pregnant dams were used for E14.5. At least two pregnant dams were necessary for these embryonic stages due to the size of the ovaries and the necessity of preparing multiple cryoblocks to maximize the chance of obtaining a section worthy of sequencing. Dissections were performed in under 20 minutes to minimize RNA degradation.

### Perinatal and adult tissue collection

All tools and dishes were first disinfected with 70% ethanol and sprayed with RNaseZap. All 1X PBS-/- was prepared using DEPC-H2O to further prevent RNase activity during dissection. For perinatal dissections i.e. postnatal (P) day 0 (day of birth) and P3, pups were anesthetized with isoflurane followed by decapitation and bisection. The lower bisected half was added to 1X DEPC-PBS -/- on ice. Ovary/oviduct complexes were quickly excised and added to a droplet of OCT. For P21 and P90, animals were euthanized using the isoflurane drop method followed by decapitation or cervical dislocation. Ovary/oviduct complexes were collected in a pool of 1X DEPC-PBS to wash off excess blood and then added to a droplet of OCT. Dissections were performed in under 10 minutes to protect RNA quality.

### Cryoblock preparation for Visium HD

Blocks were prepared with the express goal of adding as many ovaries as possible to a single mold, ideally in the same plane, to maximize the sequencing potential for each time point. To accomplish this, the ovaries that were previously added to an OCT droplet (see above) were individually removed and added side-by-side to the bottom of a 7 mm x 7 mm cryomold. See **Figure 1** for an example block that was fully loaded. Care was taken to make sure that the ovaries were in different orientations. Once the ovaries were added to the block and the space was maximally filled, OCT was carefully added slowly to the mold using a syringe, filling it to the top. If the ovaries shifted, they were carefully pushed back to the bottom with forceps avoiding the introducing of bubbles. The mold was then flash frozen using a dry ice / 100% ethanol bath. The dry ice bath was allowed to cool for at least 10 minutes before use. Flash frozen blocks were then transferred to a -80°C freezer for storage until ready to be sectioned. If there were leftover ovaries, additional blocks were made until all tissue was used.

### Cryosectioning for Visium HD

Sectioning was performed as described in 10X Genomics Visium HD Fresh Frozen Tissue Preparation Demonstrated Protocol (CG000763 Rev A). Briefly, 10-µm sections were cut using a Leica cryostat (Leica Biosystems, USA) set to -20°C. Blocks were equilibrated to the cryostat chamber temperature for at least 30 min before sectioning. Blocks were mounted with the part of the block containing the samples facing out toward the user. The cryostat blade was changed before each session, and slides were equilibrated to the chamber temperature prior to sectioning. Blocks were mounted in a mound of OCT and allowed to freeze for 2 min before mounting the chuck on the section head. Sections were cut at 10-µm intervals, and each section was captured onto a slide. To adhere a section to the slide, the flat section was gently touched with the front of a blank slide, and the OCT was melted using a finger pressed to the back of the slide. Once completely melted, the section was then added back to the cryobar and allowed to re-freeze. Two consecutive sections were placed on each slide. Slides were stored at - 80°C.

### Hematoxylin and Eosin (H&E) staining and imaging

Methanol fixation and H&E staining was performed in accordance with 10X Genomics Visium CytAssist Spatial Gene Expression for Fresh Frozen Demonstrated Protocol (CG000614 Rev A). This protocol was used because present day fixation procedures had yet to be released (see CG000763 Rev D). Following staining, slides were imaged at 20X using a Leica DM6b microscope (Leica Microsystems, USA), images were captured using a Leica K3C camera, and images were processed and stitched using LAS-X software. Downstream tissue processing was performed by the University of Colorado Anschutz Genomics Shared Resource (RRID: 021984).

### Visium HD cDNA library preparation and sequencing

Sections were hybridized overnight with 10x Genomics’ whole transcriptome probe pairs following the Visium HD Spatial Gene Expression Reagent Kits User Guide (CG000685 Rev B). After washing and ligation of bound probe pairs, the prepared slide and a Visium HD slide were processed on the CytAssist instrument to transfer and spatially barcode the ligated probes. Spatially barcoded products underwent library preparation per the Visium HD protocol. Libraries were pooled and sequenced on an Illumina NovaSeq X Plus system (150bp x 10bp x 10bp x 150bp) at the University of Colorado Genomics Shared Resource (RRID: 021984), generating approximately 500 million reads per capture area.

### Spatial transcriptomics data analysis

Loupe Browser (version 8.1.2) was used to manually outline tissue sections for the E12.5 and E14.5 datasets to exclude one testis and one adrenal gland present on those slides (see yellow asterisks in **Figure 2**). For the remaining datasets, tissue detection was performed automatically by SpaceRanger (10X Genomics, USA). Raw data were processed using SpaceRanger (version 3.1.2). SpaceRanger outputs data at the level of the 2 μm^2^ detection spots present on the Visium HD slide. Due to the sparsity of the 2 μm data, SpaceRanger also bins the 2 μm^2^ data into larger squares tiled across the capture area. Unless otherwise noted, downstream analysis was performed using 8 μm^2^ square bins, which provided the optimal tradeoff between spatial resolution, quality of cell type identification from clusters, and computational performance. Integration and clustering were performed using Seurat (*85*) (version 5.1.0). 8 μm^2^ bins with fewer than 50 reads were filtered out and data were log normalized. To facilitate efficient clustering of such a large dataset, we used sketch-based sampling to select 50,000 8^2^ μm bins for each stage while preserving rare cell types (*86*). This subset was used to perform variable feature selection, data scaling and principal component analysis (PCA). Integration of data from the different fetal and postnatal stages was performed using Seurat’s reciprocal PCA (RPCA) method. 8 μm^2^ bins were clustered using graph-based clustering (30 principal components, resolution = 0.8). Integration and clustering were projected onto the full dataset. Differential gene expression was used to identify marker genes for each cluster. De novo and known marker genes were used alongside the spatial positions and distributions of each cluster to perform cell type identification.

Although 8 μm^2^ binning performs better for clustering and cell-type annotation, the 2 μm^2^ data provide better spatial resolution for the visualization of individual genes. To reduce the sparsity of the 2 μm^2^ data while retaining the high spatial resolution, we utilized SAINSC (version 0.3.1) to perform kernel density estimation (KDE) on the 2 μm^2^ data, providing high-resolution, smoothed gene expression profiles for visualization (*48*). SAINSC was also used to assign cell types to 2 μm^2^ bins.

### Data visualization

Visualizations of expression data and clusters projected onto the tissue image were generated using Loupe Browser (10x Genomics, version 9), SpatialData (version 0.3.0) and Matplotlib (version 3.10.0) (*87*, *88*). Additional graphs were generated with ggplot2 (*89*) (version 3.5.2). Plotgardener (*90*) (version 1.14.0) was used for assembling figures.

### Single-nucleus multiome library preparation

Whole ovaries and surrounding rete ovarii tissue were collected at E16.5 and P0. Tissues were fresh frozen with dry ice in an ethanol bath, and stored at -80°C until nuclei isolation. Each timepoint included samples collected from three separate litters for a total of 39 and 33 ovaries for E16.5 and P0 respectively. Nuclei isolation was performed using a Singulator 100 instrument (S2 Genomics) per the S2 genomics Demonstrated Protocol: Nuclei Isolation from Frozen Tissue for Single Nuclei Sequencing Applications, parts A and B. The Extended Nuclei Isolation protocol was utilized. Isolated nuclei were resuspended and counted via hemocytometer, for a target recovery of 6,000 and 10,000 nuclei for E16.5 and P0 respectively. The 10x Genomics Chromium Next GEM Single Cell Multiome ATAC+Gene Expression Library Preparation Kit (16 rxns PN-1000283) was used to generate a 10x barcoded library from individual nuclei. The manufacturer’s protocol (CG000338 Rev F Chromium Next GEM Single Cell Multiome ATAC + Gene Expression) was used to generate libraries. Libraries were obtained from an estimated 5,436 and 6,268 cells per E16.5 and P0 samples respectively. Libraries were sequenced on an Illumina NovaSeq Plus instrument by Novogene (UC Davis, USA). E16.5 mean raw reads per cell were reads 602M and 682M reads for gene expression and ATAC respectively. P0 mean raw reads per cell were 664M and 730M reads for gene expression and ATAC respectively. Only the Gene Expression libraries were analyzed and use in this study.

### Single-nucleus multiome analysis

Raw single-cell multiome data from E16.5 and P0 ovaries were processed using Cellranger-arc (version 2.0.2). As this study utilizes the multiome data for directly comparing gene expression between the single-nucleus RNA-seq (snRNA-seq) with Visium HD spatial transcriptomics data, only the RNA data from the multiome were subject to downstream processing and analysis. Cellranger-arc identified 11,704 valid nuclei, 5,436 at E16.5 and 6,268 at P3. The snRNA-seq count matrices were processed using decontX (version 1.8.0) to correct for the potential effect of ambient RNA (*91*). To identify and remove potential doublets generated during droplet encapsulation, scDblFinder (version 1.24.0) (*92*) was used to identify an appropriate, data-driven threshold for removing probable doublets. Nucleus barcodes with an scDblFinder doublet score greater than 0.77 (E16.5) or 0.76 (P0) were filtered out. Additional quality control filtering of nucleus barcodes was performed to remove nuclei meeting the following criteria: more than 5 median absolute deviations (MADs) above or below the median UMIs; more than 5 MADs below the median genes per nucleus; greater than 1% of counts from mitochondrial genes. An additional 160 barcodes were removed based on preliminary analysis indicating that these barcodes derived from dead cells. Altogether, a total of 10,241 nuclei were retained for downstream analysis, 4,751 from E16.5, and 5,490 from P0. Integration and clustering were performed using Seurat. First, filtered snRNA-seq data were subject to variable feature selection, data scaling, and PCA. Integration across time points was performed using Seurat’s RPCA integration. Graph-based clustering was performed at a resolution of 0.8 using the top 50 principal components, and a UMAP embedding was computed for visualization. Marker genes were identified and used to annotate cell types.

### Spot deconvolution and differential expression for spatial transcriptomic data

Spot deconvolution was performed to estimate the proportion of RNA contributed from different cell types in each 8 µm bin of the Visium HD spatial transcriptomic data. Deconvolution was performed using robust decomposition of cell type mixtures (RCTD) as implemented through the spacexr R package (version 1.2.0). The snRNA-seq data from E16.5 and P0 were used to perform deconvolution separately for the spatial transcriptomes from E16, E18, P0, and P3. RCTD was run in doublet mode to indicate that each 8 µm bin is likely to span no more than two separate cell types. From the RCTD results, we extracted the cell type from the reference snRNA-seq dataset predicted to contribute the most RNA to that spot, as well as the relative proportion of RNA contributed by additional cell types in the reference dataset. These results allowed us to filter the 8 µm bins in the spatial dataset to include only those likely to contain RNA from a single cell type (**Fig. S4**). Using the results of the spot deconvolution, we performed differential expression analysis to compare granulosa cells at the center of the ovary to those at the surface at E16.5, E18.5, and P3, while removing any spots that have contaminating RNAs from non-granulosa cell types. First, we classified granulosa cell types based on their regional patterns within the ovary. 8 µm bins annotated as active granulosa or IOR-pregranulosa mixture were classified as being enriched at the center of the ovary. Bins annotated as early granulosa or oocyte-granulosa mix were classified as being enriched at the ovarian surface. These spots were further filtered based on the RCTD deconvolution results. Only bins that were predicted to have 80% of the RNA contributed by a granulosa cell type were retained for differential expression analysis. Next, we took advantage of the high number of ovaries present in each slide by performing pseudobulk differential expression analysis in which we pooled all data from spots classified either as surface or center in each individual ovary on each slide. The pseudobulk approach allowed us to perform differential expression analysis between center and surface granulosa cells with 6-7 replicates at each stage. Following pseudobulk aggregation, differential expression was determined using standard DESeq2 analysis (*50*).

### Whole-Mount Immunofluorescence

Pregnant and perinatal female CD-1 mice were euthanized according to approved institutional regulations. Fetal and perinatal ovaries were dissected into 1X phosphate-buffered saline (PBS). Following dissection, ovary samples were fixed in 4% paraformaldehyde (PFA)/PBS for 30 minutes, rocking at room temperature (RT). Samples were then rinsed in 1X PBS followed by a wash in 1X PBS for 30 minutes, rocking at RT. Subsequently, ovaries were dehydrated in an increasing methanol (MeOH)/PBS gradient: 25%, 50%, 75% and 100% MeOH each for 15 minutes rocking at RT. Samples were then stored in 100% MeOH at -20°C until staining. For staining, samples were rehydrated via a decreasing MeOH/PBS gradient: 75%, 50% and 25%, followed by a wash in 1X PBS for 15 minutes rocking at RT. Samples were then permeabilized for 20 minutes in a PBS 0.1% Tx-100 solution rocking at RT. Following permeabilization, samples were incubated in freshly prepared block solution (PBS 1% Tx-100; 10% horse serum) rocking at RT for 1 hour. During this incubation, perinatal ovaries were carefully removed from the ovarian capsule using fine forceps, allowing for optimal ovarian permeability. After blocking, samples were incubated with primary antibodies overnight in at 4°C without rocking. Primary antibodies were diluted according to Supplementary Table 1 in 250uL block solution. The next day, samples were washed three times for 20 minutes each with PBS 0.1% Tx-100 rocking at RT. Samples were then incubated with secondary antibodies (**Supplemental Table 1**) diluted at 1:500 in 250 µL block solution. The 2°Ab solution was filtered using a syringe driven 22 µm filter before adding it to the samples. Samples were then placed in a dark box and incubated in filtered 2°Ab solution overnight at 4°C without rocking. Following incubation, samples were washed three times for 10 minutes each with PBS 0.1% Tx-100 rocking at room temperature. Samples were immediately mounted for imaging using CUBIC R2 agarose as previously described (*93*).

### RNAscope assay

Females were euthanized according to approved institutional regulations. Ovaries were dissected into 1X PBS and fixed in 4% PFA/PBS overnight at 4°C. Following fixation, samples were washed three times 15 minutes with 1X PBS rocking at RT and subsequently transferred into 30% sucrose/PBS overnight at 4°C. Samples were then prepared for embedding. Specifically, ovaries were removed from the 30% sucrose/PBS solution using forceps and dipped twice into OCT embedding compound. Samples were then transferred into disposable plastic cryomolds containing a layer of frozen OCT. A second layer of OCT was used to cover the sample(s) and the cryomold was placed over dry ice to freeze. Sample blocks were then stored at -80°C until used for cryosectioning. Slides for staining were allowed to acclimate to -20°C for 30 minutes to an hour before cryosectioning. Blocks containing ovary samples were mounted onto the cryostat chuck using OCT and the sample(s) were sectioned into 10μm thick slices using a Leica CM 1950 Cryostat (Leica Biosystems). Sample sections were mounted directly onto Superfrost Plus Microscope Slides (Fisherbrand) and stored at -80°C for up to three months before use. Sample pre-treatment for fixed-frozen tissue was performed per the protocol outlined in the RNAscope Multiplex Fluorescent Reagent Kit v2 User Manual UM 323100/Rev C/Effective Date: 02/18/2025 pages 44 – 47 (ACDBio, Inc., USA). Target retrieval was not performed. Following dehydration, a hydrophobic barrier was applied and allowed to fully dry before proceeding with hydrogen peroxide and protease treatment. The RNAscope assay (probe hybridization – HRP-C1 signal) was performed per Chapter 6 of the RNAscope Multiplex Fluorescent Reagent Kit v2 User Manual UM 323100/Rev C/Effective Date: 02/18/2025. Probes used are listed in **Supplemental Table 2**. TSA Vivid 650 dye was used at a 1:1000 dilution in TSA buffer. Slides were counterstained with DAPI (provided with RNAscope kit) and mounted in Dako Fluorescence Mounting Medium (Agilent Ref: S3023).

### Image acquisition

All ovary samples that underwent RNAscope and whole-mount IF were imaged on an Oxford Instruments Andor Dragonfly 202 Spinning Disk Confocal microscope with a 40μm disk pinhole (Oxford Instruments, Abingdon, UK), with an Applied Scientific Instrumentation Piezo stage controller (Applied Scientific Instrumentation, OR, USA). Images were acquired using a 25X water immersion (NA 0.95) objective (Leica Microsystems, Wetzlar, Germany). Images were captured by an Andor Zyla 4.2 plus cMOS camera (Oxford Instruments Andor, Abingdon, UK). We used a high-power laser engine (HLE) equipped with 405, 637 and 561 nm lasers to visualize nuclei (DAPI and Hoechst dyes), Alexa Fluor dye 647, and Cy3. Confocal ZX-stacks were captured with a Z-interval of 1 µm. Image panels and figures were assembled in Adobe Photoshop and each image and channel was individually adjusted for exposure, brightness and contrast for optimal display of fluorescence signals.

### Data and code availability

Sequencing data have been deposited in the Gene Expression Omnibus (GEO) with accession # GSE307688. All analysis code is available on GitHub: https://github.com/McKeyLab/Ovary_VisiumHD.

## Acknowledgments

This study was supported by start-up funds from the CU Anschutz Medical Campus, School of Medicine, Department of Pediatrics, Section of Developmental Biology to JM, by the Boettcher Foundation Webb-Waring Biomedical Research Award to JM and by funding from the National Institutes of Health to JM (grant #R00HD103778). This study was supported in part by the University of Colorado Anschutz Medical Campus Genomics Shared Resource (RRID: 021984). We are grateful to all members of the McKey laboratory and the Roberson laboratory for helpful discussions and suggestions on the work presented here.

## Author contributions

Conceptualization: ASM, TJG, JM,

Methodology: ASM, TJG, CD, JJ, JM,

Investigation: ASM, TJG, CD, JJ, JM

Visualization: ASM, TJG, CD, JJ, JM

Supervision: JM

Writing—original draft: ASM, TJG, JM

Writing—review & editing: ASM, TJG, CD, JJ, JM

## Competing interests

All other authors declare they have no competing interests.

## Data and code availability

Sequencing data have been deposited in the Gene Expression Omnibus (GEO) and will be made public upon peer-reviewed publication and shared upon reasonable request in the meantime. All analysis code is available on GitHub: https://github.com/McKeyLab/Ovary_VisiumHD.

## Notes

### Competing Interest Statement

The authors have declared no competing interest.

### Summary of Updates

Most of the manuscript was rewritten to include new results. Two new authors were added to reflect contribution to new results.

