## Supplemental Materials for "Single-cell-scale spatial transcriptome reveals early regional priming of the developing mouse ovary"

Anthony S. Martinez *et al.*

**This PDF file includes:**

Supplementary Text
Figs. S1 to S8
Tables S1 to S2

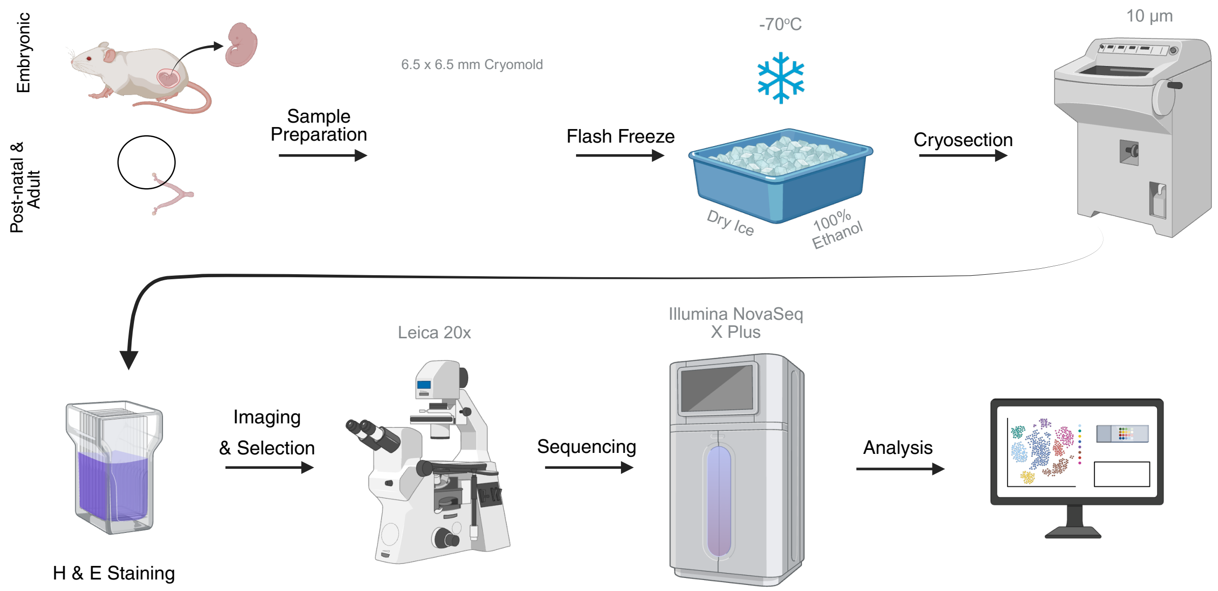


**Fig. S1.**

**Experimental design**. Fetal, perinatal, and adult ovaries were dissected, embedded in a 6.5 x 6.5 mm cryomolds, flash frozen, and cryosectioned. 10 µm sections were stained with H & E, imaged, and processed for HD Visium. RNA was then captured using CytAssist followed by cDNA library preparation and sequencing. Data were processed, analyzed and projected onto the H&E images using 8 $\mu m^2$ bin resolution. Created in BioRender. Martinez, A. (2025) https://BioRender.com/sytaju5

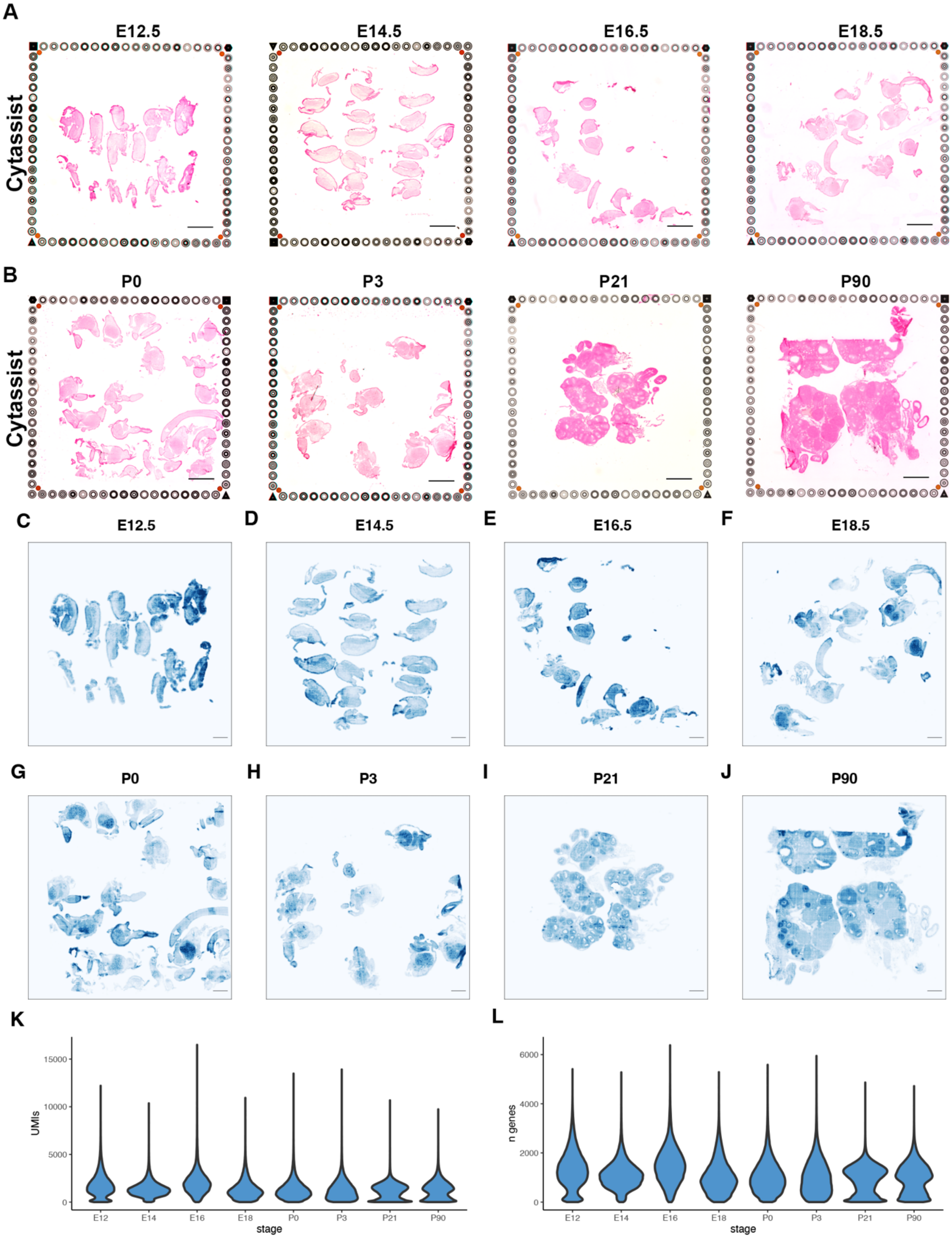
A
E12.5
E14.5
E16.5
E18.5
Cytassist
B
P0
P3
P21
P90
Cytassist
C
E12.5
D
E14.5
E
E16.5
F
E18.5
G
P0
H
P3
I
P21
J
P90
K
UMIs
15000
10000
5000
0
E12
E14
E16
E18
P0
P3
P21
P90
stage
L
n genes
6000
4000
2000
0
E12
E14
E16
E18
P0
P3
P21
P90
stage

**Fig. S2.**

**Technical validation of UMI complexity and gene capture**. (**A**) CytAssist images obtained prior to sequencing and subsequently used for alignment of capture areas to regions of tissue. Scale bars, 1000 µm. (**B**) UMI counts for each timepoint projected onto histology images of the sequenced tissue. Scale bars, 500 µm. (**C**) Violin plot showing distribution of UMI counts per capture area for each timepoint. (**D**) Violin plot showing the distribution of the number of genes detected per capture area at each stage.

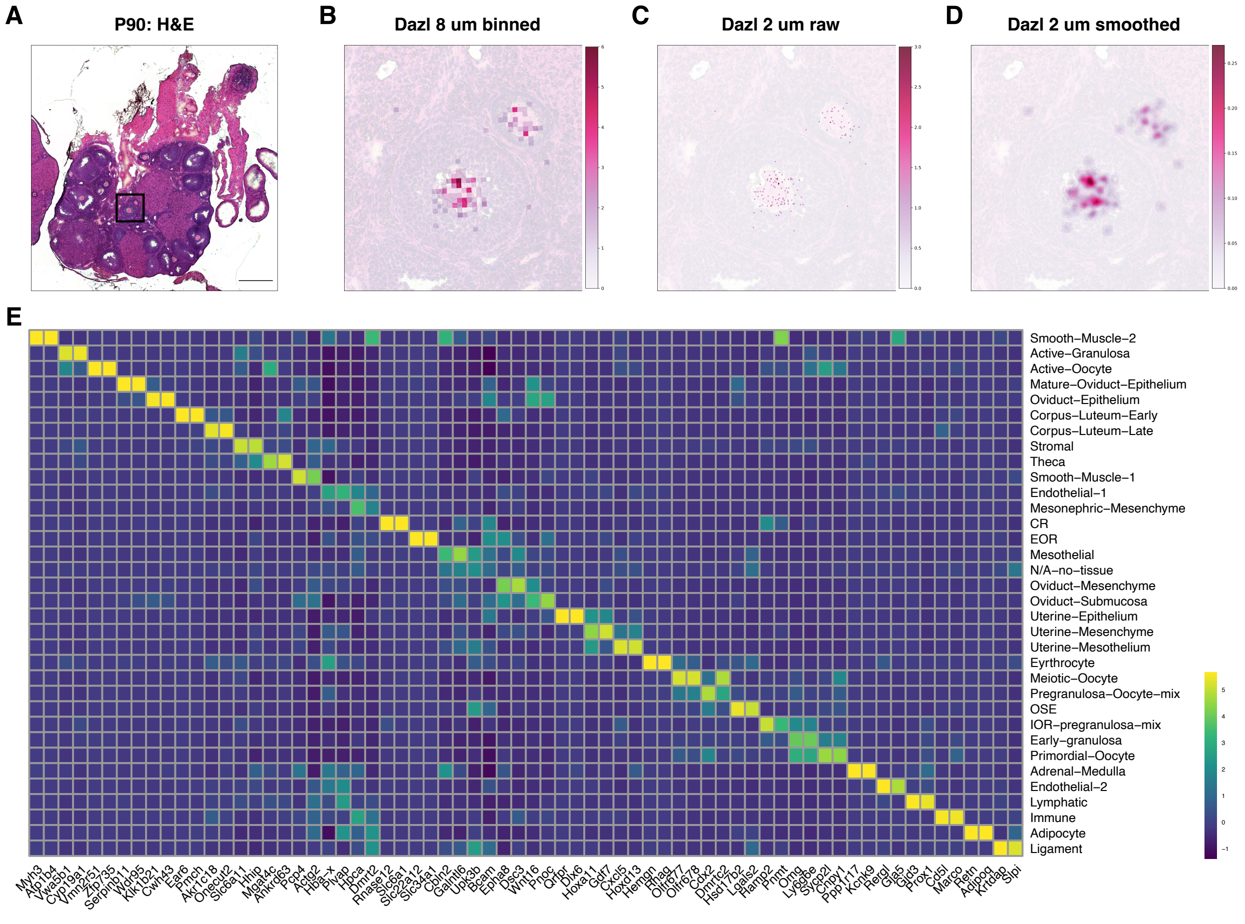


**Fig. S3.**

**SAINSC based smoothing validation and novel uniquely expressed genes of integrated clusters.** (**A - D**) Comparison of the spatial resolution between 8 μm$^2$ bins and SAINSC smoothed 2 μm$^2$ bins on a P90 ovary to display expression of the oocyte marker Dazl. (**E**) Heatmap displaying the 34 integrated clusters along with their top two uniquely expressed genes. Yellow indicates higher expression; blue indicates lower expression.

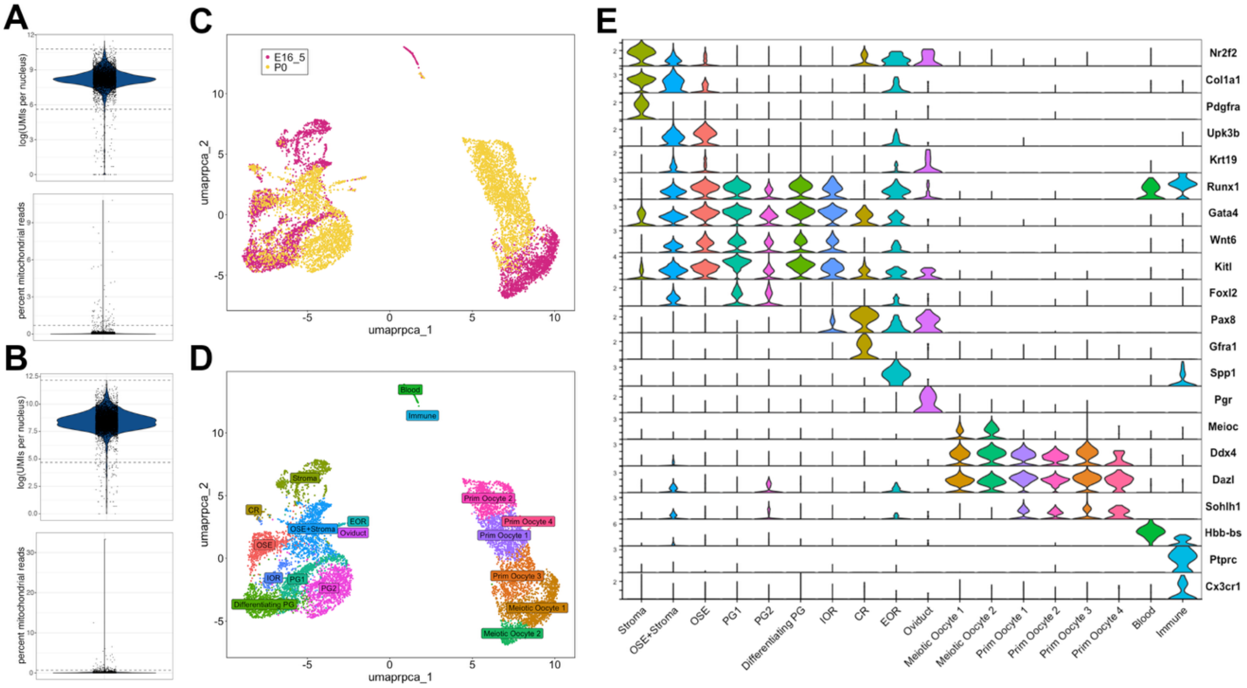


**Fig. S4.**

**QC and cluster annotations for SnRNAseq of E16.5 and P0 ovary-rete ovarii complexes.** (**A – B**) Violin plots illustrating sample quality control for snRNAseq data at E16.5 (**A**) and Postnatal day (P)0 (**B**). Top rows show log(UMIs) per nucleus as a measure of library complexity, bottom rows show percent mitochondrial reads, as a measure of cell lysis and cytoplasmic RNA contamination. Dotted lines represent data filtering thresholds for further analysis. (**C**) Seurat UMAP embedding showing analysis of the rpca-integrated single-nucleus dataset from E16.5 (*cherry*) and P0 (*yellow*) samples. (**D**) Seurat UMAP embedding showing cluster analysis of the single-nucleus rpca-integrated dataset using resolution = 0.8. 18 clusters were identified and annotated using known gene markers and Gene Ontology analysis. (**e**) Seurat Stacked Violin plot showing expression of marker genes in each cluster of the integrated dataset. True correspond to filtered bins and False correspond to unfiltered bins.

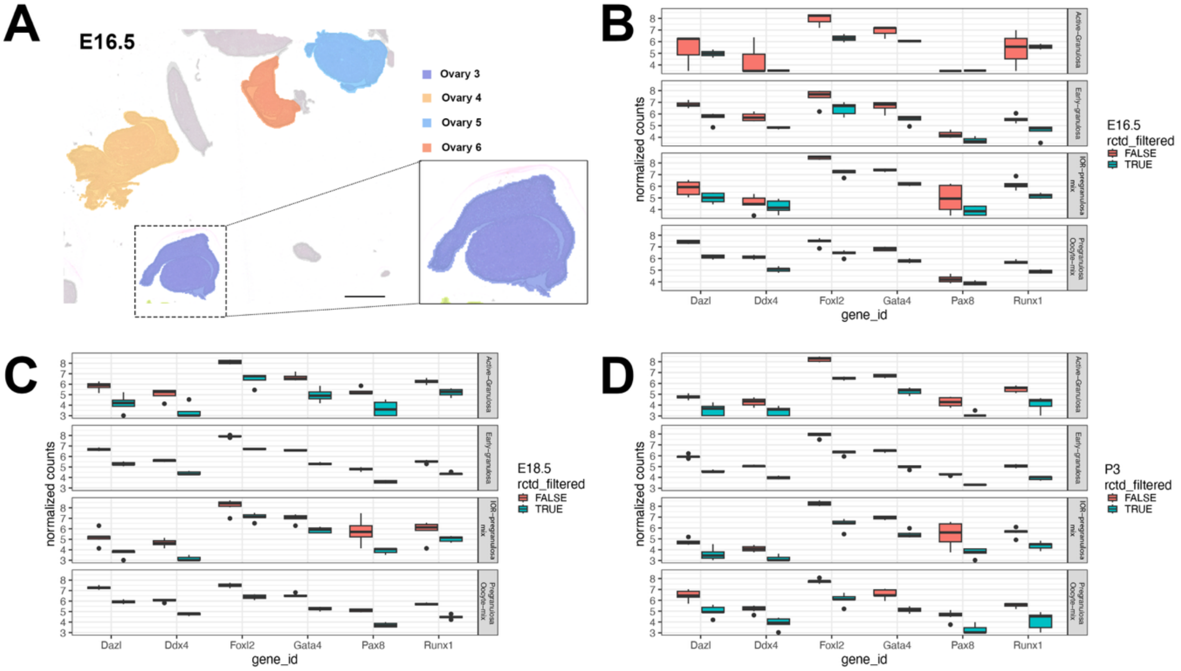


**Fig. S5.**

**Pseudobulking and deconvolution QC on E16.5, E18.5 and P3 mouse ovaries.** (**A**) Example of ovary replicate selection for psudobulk based analysis at E16.5. Ovaries containing sections through the cortex and medulla were sequentially outlined in Loupe Browser (Version 8.0.0). The bins associated with each replicate ovary were exported in a csv and used for downstream analysis. Ovaries 3 – 6 are four of the six replicates used at E16.5. Scale bar, 500 µm. (**B – D**) Box plots of normalized counts for *Dazl, Ddx4, Foxl2, Gata4, Pax8,* and *Runx1* for each of the granulosa clusters selected for analysis (*Active Granulosa, Early Granulosa, IOR-Pregranulosa Mix,* and *Pregranulosa-Oocyte Mixed*) at E16.5, E18.5 and P3 after RCTD based filtering. "False" demark unfiltered bins and "True" demark filtered bins.

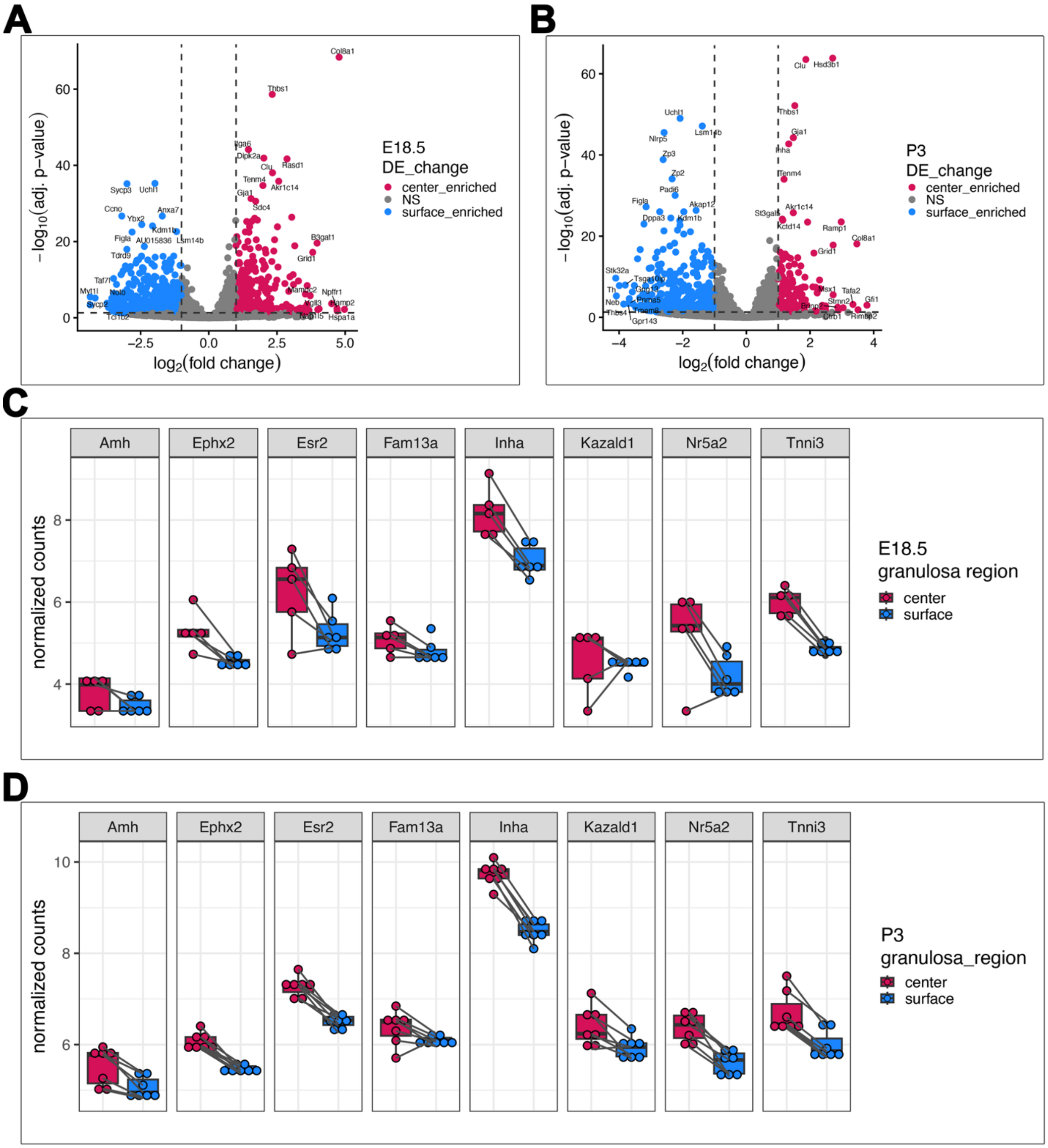


**Fig. S6.**

**Differential gene expression between center and surface PGs at E18.5 and P3.** (**A – B**) Volcano plots of differential gene expression between center enriched (*active granulosa*, and *IOR-pregranulosa mixed* clusters) and surface enriched (*oocyte-granulosa mix* and *early granulosa* clusters) 8 μm$^2$ bins at E18.5 (n = 6) and P3 (n = 7). Bins were deconvolved and pseudobulked prior to analysis for ≥80% granulosa identity. (**C – D**) Box plot of normalized expression (varience-stabilized transformation) from active signature genes between the center and surface at E18.5 and P3: *Amh, Ephx2, Esr2, Fam13a, Inha, Kazald1, Nr5a2,* and *Tnni3*. Boxplots are overlaid with dotplots showing values for individual ovaries. Lines connect values for the center and surface of the same individual ovary.

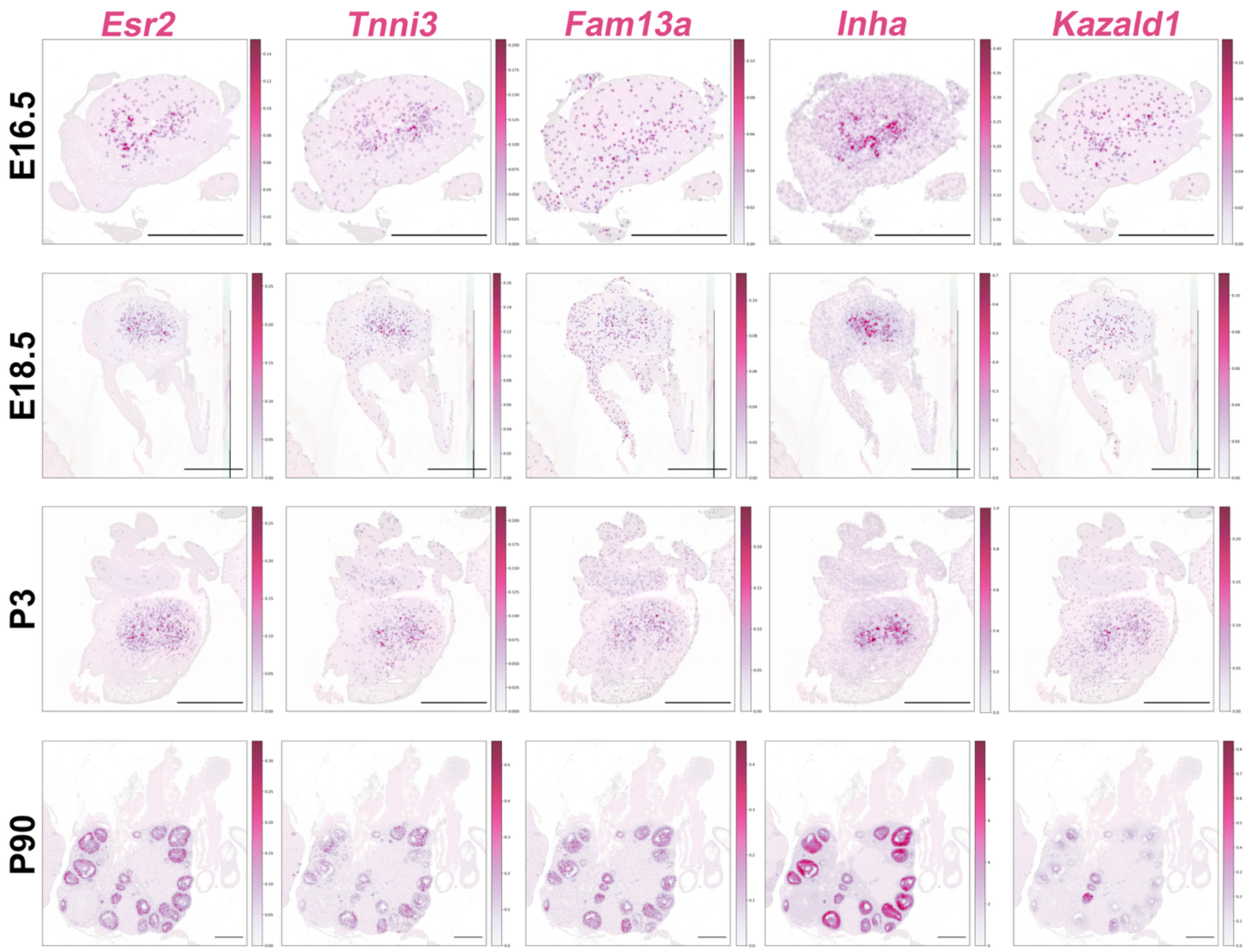


**Fig. S7.**

**Spatial gene expression of active signature genes in fetal and postnatal ovaries.** (**B**) Smoothed 2 μm$^2$ spatial expression of *Esr2, Tnni3, Fam13a, Inha,* and *Kazald1* projected onto a E16.5, E18.5, P3 and P90 ovaries. Scale bar, 500 μm.

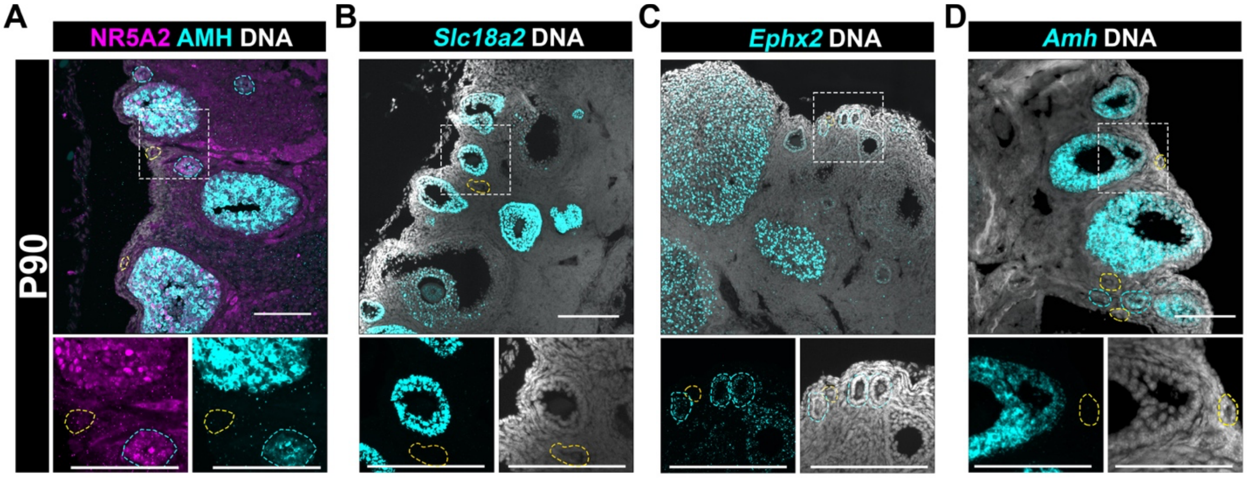


**Fig. S8.**

**Validation of active signature genes in growing follicles using immunofluorescence and RNAscope.** (**A**) Maximum intensity projection (MIP) from confocal Z-stacks of ovaries collected at P90, cryosectioned at 10 µm and immunostained for NR5A2 (*magenta*) and AMH (*cyan*), and counterstained with Hoechst nuclear dye (DNA, *grayscale*). Primordial follicles outlined in yellow, primary follicles outlined in cyan. Scale bar, 100 µm. (**B-D**) MIPs from confocal Z-stacks of ovaries collected at P90, cryosectioned at 10 µm and processed for RNAscope using probes against mouse *Slc18a2* (**B**), mouse *Ephx2* (**C**), and mouse *Amh* (**D**)and counterstained with DAPI nuclear dye (DNA, *grayscale*). Images in the bottom rows are split-channel zoomed-in insets from the dotted rectangles in top row images. Yellow outlines indicate quiescent primordial follicles, while cyan outlines indicate growing primary follicles. Scale bars, 100 µm.

**Table S1.**

| Primary Antibody | Host Species | Dilution | Supplier | Catalog Number | RRID | Secondary Antibody | Dilution | Supplier | Catalog Number | RRID |
|---|---|---|---|---|---|---|---|---|---|---|
| AMH | Rabbit | 1:250 | abcam | ab272221 | AB_3738533 | Alexa Fluor 647 Donkey Anti-Rabbit | 1:500 | Jackson ImmunoResearch | 711605152 | AB_2492288 |
| RUNX1 | Rabbit | 1:500 | abcam | ab92336 | AB_2049267 | | | | | |
| AF488-SYCP3 | Mouse | 1:250 | abcam | ab205846 | AB_10678841 | N/A (Conj. primary) | | | | |
| NR5A2 | Goat | 1:250 | R&D Systems | AF2030 | AB_2277369 | Cy3 anti-goat | 1:500 | Jackson ImmunoResearch | 705165147 | AB_2307351 |
| FOXL2 | Goat | 1:250 | abcam | ab5096 | AB_304750 | | | | | |

Supplemental Table 1. List of antibodies used in this study

**Table S2.**

| Target | Probe cat # (ACDBio) | Chemistry | Detection channel | Accession # |
|---|---|---|---|---|
| *mm Ephx2* | 558701 | C3 | 650 | NM_001271421.1 |
| *mm Slc18a2* | 425331 | C3 | 650 | NM_172523.3 |
| *mm Amh* | 489811 | C1 | 650 | NM_007445.2 |

Supplemental Table 2. List of RNAscope probes used in this study
